# A single-cell based precision medicine approach using glioblastoma patient-specific models

**DOI:** 10.1101/2021.01.13.426485

**Authors:** James H. Park, Abdullah H. Feroze, Samuel N. Emerson, Anca B. Mihalas, C. Dirk Keene, Patrick J. Cimino, Adrian Lopez Garcia de Lomana, Kavya Kannan, Wei-Ju Wu, Serdar Turkarslan, Nitin S. Baliga, Anoop P. Patel

**Affiliations:** Institute for Systems Biology, Seattle, WA; Department of Neurological Surgery, University of Washington, Seattle, WA; Human Biology Division, Fred Hutchinson Cancer Research Center, Seattle, WA; Department of Pathology, University of Washington, Seattle, WA; Center for Systems Biology, University of Iceland, Reykjavik, Iceland; Departments of Microbiology, Biology, and Molecular Engineering Sciences, University of Washington, Seattle, WA; Brotman-Baty Institute for Precision Medicine, University of Washington, Seattle, WA

## Abstract

Glioblastoma (GBM) is a heterogeneous tumor made up of cell states that evolve over time. Here, we modeled tumor evolutionary trajectories during standard-of-care treatment using multimodal single-cell analysis of a primary tumor sample, corresponding mouse xenografts subjected to standard of care therapy, and recurrent tumor at autopsy. We mined the multimodal data with single cell SYstems Genetics Network AnaLysis (scSYGNAL) to identify a network of 52 regulators that mediate treatment-induced shifts in xenograft tumor-cell states that were also reflected in recurrence. By integrating scSYGNAL-derived regulatory network information with transcription factor accessibility deviations derived from single-cell ATAC-seq data, we developed consensus networks that regulate subpopulations of primary and recurrent tumor cells. Finally, by matching targeted therapies to active regulatory networks underlying tumor evolutionary trajectories, we provide a framework for applying single-cell-based precision medicine approaches in a concurrent, neo-adjuvant, or recurrent setting.

**Summary:** Inference of mechanistic drivers of therapy-induced evolution of glioblastoma at single cell resolution using RNA-seq and ATAC-seq from patient samples and model systems undergoing standard-of-care treatment informs strategy for identification of tumor evolutionary trajectories and possible cell state-directed therapeutics.

## Introduction

GBM is a highly lethal malignancy of the brain that is refractory to standard-of-care (SOC) therapy, which consists of surgery, radiation (XRT), and chemotherapy with the DNA-alkylating agent, temozolomide (TMZ) (*1*–*3*). Despite aggressive treatment, median survival is only 14-17 months. Previous studies have shown that GBM tumors are made up of a complex ecosystem of normal cell types and malignant tumor cell states (*4*–*6*). Given this intratumoral heterogeneity, a multimodal systems biology approach is well-suited to characterize and uncover the mechanistic drivers of genetic and epigenetic programs that distinguish cell states within the tumor. Moreover, non-genetic, treatment-induced shifts in cell state occur along trajectories that are as yet unknown.

Here, we develop a framework to model and characterize non-genetic cell states that comprise a GBM tumor at the outset of the disease and during treatment-induced evolution (Fig. 1A). This framework, applied to an individual patient, is based on single-cell multi-omic analysis (scRNA-seq, scATAC-seq) of the initial patient biopsy, a time series of SOC-treated patient-derived xenografts (scRNA-seq), and of the recurrent tumor treated with XRT at autopsy (scRNA-seq, scATAC-seq). From our analysis, we identified multiple mechanistic drivers of treatment-induced transitions in the epigenetic states of tumor cells (*6*–*8*). As proof of concept, we then identified potential therapeutics that could target specific cell states during various stages of tumor evolution. This work collectively provides a framework to model tumor evolution during treatment and implement systems biology approach-based single-cell analysis for the rational design of precision therapeutic regimens.

**Figure 1.**
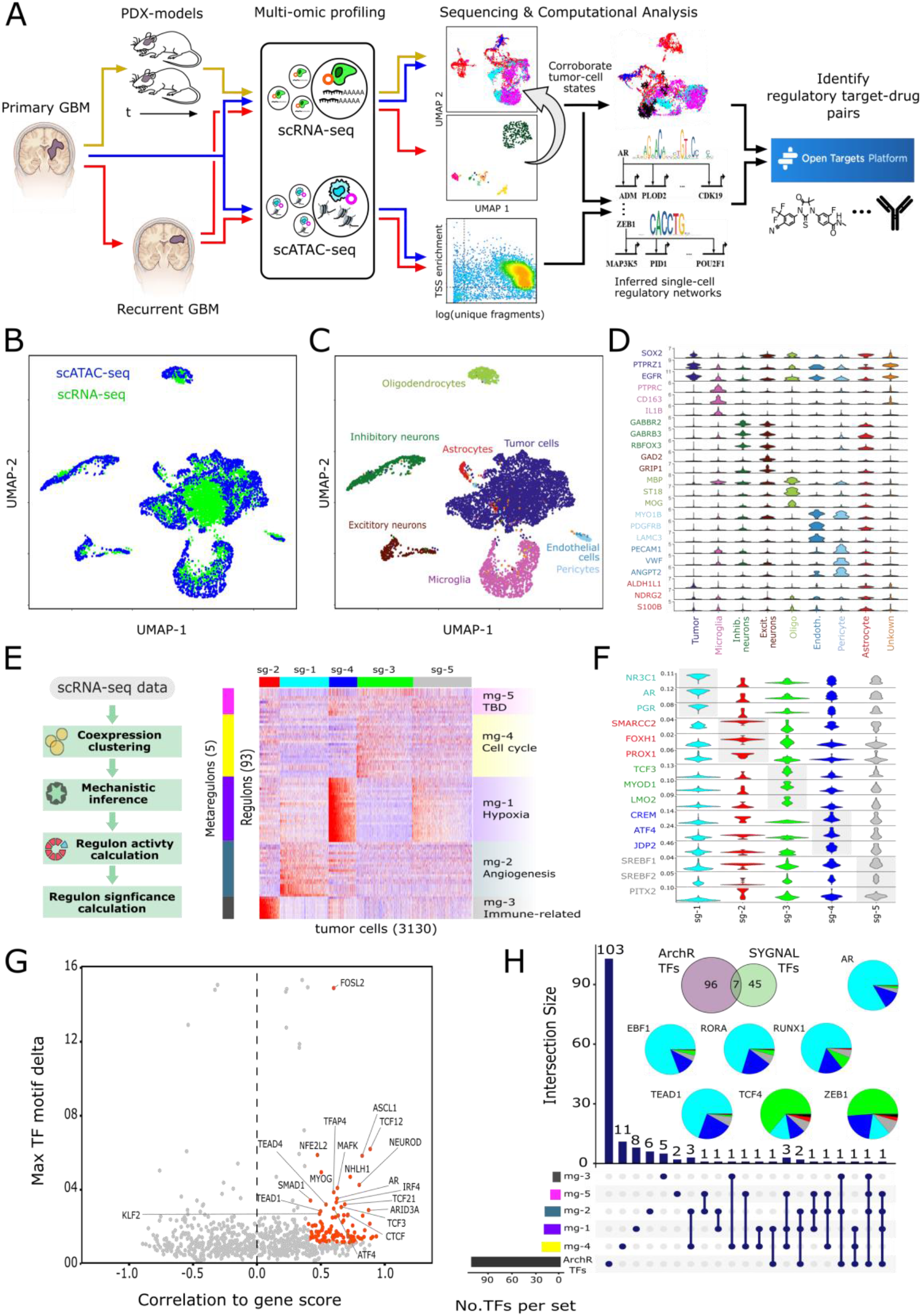
Multi-modal single-cell characterization of UW7 primary tumor biopsy. **(A)** Schema of overall proof-of-concept modeling and analytical framework used to characterize intratumoral heterogeneity using multi-omic, single-cell level analysis and drive precision care for individual GBM patient. **(B)** UMAP plots of integrated scATAC-seq and scRNA-seq profiles. **(C)** Cell-type annotation of integrated single-cell data based on established cell-type-specific gee sets. **(D)** Violin plots of cell-type marker gene expression (log2[normalized counts + 1]). **(E)** scSYGNAL/MINER analysis of scRNA-seq profile of cells from UW7 primary tumor biopsy. Heatmap indicates activity (z-scores) of co-regulated gene modules, (i.e., regulons [rows] across cells [columns]). Groups of regulons sharing similar activity patterns across cells define transcriptional programs (mg-X), each enriched with specific biological functions. Groups of cells sharing similar activity profiles across regulons define transcriptional network states (sg-X). **(F)** Violin plots showing distribution of standardized deviation accessibility scores (deviation scores) of top three TF binding motifs per scATAC-seq sample. **(G)** Scatter plot showing positive TF regulators (orange dots) per scATAC-seq profiles. Positive TF regulators have deviation scores that correlate with their corresponding inferred gene expression (gene score) values (correlation ≥ 0.4, FDR-adjusted p-value ≤ 0.1) and have a maximum inter-sample group deviation score difference in the upper 50% quantile. The top 20 TFs having maximal radial distance from the origin are labeled for reference. **(H)** Upset plot delineates the number of TFs identified in single-cell samples via scSYGNAL and ArchR analysis and the number of shared TFs across all combinations of TF sets associated with transcriptional program-based groups and ArchR TF sets. SYGNAL/MINER analysis of scRNA-seq profiles and ArchR analysis of scATAC-seq profiles identified 52 and 103 “active” TFs, respectively, and resulted in seven consensus TFs. Pie charts indicate the composition of single cells from each transcriptional network state associated with a positive deviation score for each of the consensus TFs.

## Results

### scRNA-seq and scATAC-seq analysis reveals multiple transcriptional network states in glioblastoma

We first aimed to create a reference landscape of cell types and tumor cell states by integrating both scRNA-seq and scATAC-seq data into the same latent space using previously established methods (Fig. 1B) (*9*). We then identified marker genes for each major cluster and manually curated cell-type annotations (Fig. 1C, 1D). Based on the curated cell-type annotation, we analyzed a total of 7,723 single cells from the primary tumor, which consisted of 358 (4.6%) oligodendrocytes, 1,169 (15.1%) microglia, 134 (4.2%) pericytes/endothelial cells, 305 (3.9%) excitatory neurons, 568 (7.4%) inhibitory neurons, and 103 (1.3%) astrocytes. Further, we identified 4,924 tumor cells, which made up 63.7% of single cells collected from the primary sample, based on a characteristic gain of chromosome 7 and loss of chromosome 10 in GBM (*10*) (Fig. 1B, 1C, S1). The identities of subpopulations were confirmed by statistically significant differential expression of marker genes including *SOX2, PTPRC. MBP, RBFOX3, VWF*, and *PTPRZ1* for various cell subtypes (Table S1).

To identify regulator networks in scRNA-seq data, we adapted the Systems Genetics Network AnaLysis (SYGNAL) platform (*11*) to analyze the scRNA-seq profiles (scSYGNAL) and identify distinct epigenetic programs that were differentially active and distinguished subpopulations of tumor cells in the primary tumor. Briefly, using biclustering of scRNA-seq data, we inferred regulons (i.e., sets of genes sharing similar expression patterns and putatively co-regulated by the same transcription factor (TF) or miRNA across a sub-population of single cells). Our analysis revealed the mechanistic co-regulation of 809 genes across 160 regulons by at least 65 TFs and 141 miRNAs. We then used the Mining for Node Edge Relationships (MINER) algorithm (*12*) to identify a subset of 93 significant regulons that included 52 unique TFs regulating 454 target genes (Methods, Tables S2 and S3). These 93 regulons were further clustered into 5 metaregulon groups (i.e., transcriptional programs) with distinct regulon activity profiles across the 3,130 tumor cells analyzed with scRNA-seq (Fig. 1D, E). Moreover, functional enrichment analysis (*13*), revealed distinct biological processes and molecular functions associated with these programs (Fig. 1D, Tables S4-S8), the amalgamation of which we defined as a transcriptional network state.

### scSYGNAL identifies a network of regulators mediating treatment-induced shifts in xenograft tumor-cell states also reflected in recurrent disease

Based on these transcriptional network states, we clustered the tumor cells into 5 groups of cells sharing similar network states, with each group exhibiting upregulated activity of specific programs. For example, the SG-4 subpopulation expressed increased activity in program 1, enriched for hypoxia-associated genes like *VEGFA, PLOD2*, and *PDK1*. In addition, the SG-4 subpopulation exhibited decreased activity in cell-cycle-related regulons (transcriptional program 4), suggesting that this group is comprised mainly of non-proliferative cells. Further, statistical analysis revealed an over-enrichment of non-proliferating cells in SG-4 (Methods, Table S9) (*14*). The SG-1 subpopulation exhibited high activity for program 2, which includes an enrichment of angiogenesis-related genes, suggesting that this subpopulation may be involved in tumor neovascularization. The SG-3 group consists of tumor cells that expressed increased activity in regulons associated with the G2M checkpoint and E2F targets, mirroring gene expression behavior of proliferative GBM cells (*15, 16*). Similarly, proliferating cells were over-enriched in the SG-3 population (Table S9). Conversely, the SG-5 subpopulation did not exhibit distinct upregulation of any one distinct program. Rather, these cells expressed faint signatures of multiple regulons across multiple programs. It is possible that this subpopulation represents tumor cells transitioning between states or cells that are primed for expression of a variety of programs.

To place these results into a broader context, we compared the 52 unique TFs for the 93 significant regulons to those TFs deemed to be essential for GBM-specific proliferation via a genome-wide CRISPR-Cas9 screen (*17*). Of the 52 unique TFs identified in the MINER analysis of the scRNA-seq data, 17 (33%, *p* = 0.072) overlapped with the TFs essential for GBM tumor cell proliferation, indicating that our single-cell based network analysis identified relevant GBM-specific regulatory mechanisms (Table S10).

In an independent and orthogonal approach, we also analyzed scATAC-seq data to identify TF motifs that were enriched in differentially accessible regions of the genome across the tumor cell population. Using the ArchR package and chromVAR method (*18, 19*), we identified 103 TFs as positive regulators, i.e., those TFs having significant motif deviation scores that correlate with their inferred gene expression values (Fig 1F). By comparing the set of TFs identified from ArchR analysis of scATAC-seq data and the set from the SYGNAL/MINER analysis of scRNA-seq data, we identified a consensus set of 7 TFs (*AR, TEAD1, RUNX1, RORA, EBF1, ZEB1*, and *TCF4*). Notably, only a subset of these genes showed significant differential expression across the tumor population when examining their expression levels in transcriptional data, implying that many of these regulators would not have been identified using standard scRNA-seq data analyses, such as shared nearest neighbor-based cluster identification (Fig. S2) and subsequent differential expression (Table S11). Interestingly, this set of TFs has potentially important roles in tumor biology. Extensive data implicate the role of *TEAD1* in YAP/Hippo signaling in gliomagenesis and other neoplastic processes (*20*–*23*). *TCF4* is a mediator of Wnt/β-catenin signaling and correlates with glioma progression via effects on AKT2 (*24*). Similarly, *ZEB1* is a key regulator in epithelial-to-mesenchymal (proneural-to-mesenchymal) transition and associated chemoresistance mechanisms (*25, 26*). *AR* signaling has been shown to be active in GBM cells *in vitro* and may relate to radiation resistance (*27, 28*). Based on our analysis of integrated transcriptional and chromatin accessibility data, we conclude that these TFs likely play key roles in the regulation of tumor cell states in the primary tumor.

Having assessed the regulatory networks of cell states within the primary tumor, we then sought to understand how tumor cell states change over time. Tumors evolve due to both intrinsic pressures (i.e., acquired mutations, tumor-microenvironment interactions) and extrinsic pressures (therapeutic intervention). In particular, we were interested to see whether treatment-induced evolution of tumor cell-states resulted in selection of preexisting states, induction of novel states, or a combination of both. To address these questions, we modeled treatment-induced cell-state changes by applying SOC therapy (XRT/TMZ) to a cohort of patient-derived xenografts (PDX) created from the patient’s tumor. Tumor cells isolated directly from the patient were injected orthotopically into immunocompromised mice without any intervening culture. We obtained scRNA-seq data on samples collected from SOC-treated xenografts at 24 and 72 hours after completion of therapy and corresponding untreated controls. We then performed batch integration and dimensionality reduction of the scRNA-seq data to visualize, identify, and compare cell states of xenograft samples to those of the corresponding primary tumor (Fig 2A). Interestingly, ∼33% of the primary tumor cells in the UMAP embedding had a nearest neighbor originating from an untreated PDX mouse (Supplemental Fig. S3). Closer examination of the primary tumor cells having an untreated-PDX cell as a nearest neighbor revealed that these tumor cells represented all SG groups, suggesting that the PDX models were able to reproduce the majority of the cell states observed in the primary tumor. Similarly, statistical analysis of tumor cell sources (i.e., primary vs. treated PDX) within clusters identified via shared nearest neighbor (SNN) modularity optimization (*29*) revealed that several SNN clusters were enriched for tumor cells from both the primary tumor and PDX mice (Fig. S2). Taken together, these results support the idea that the transcriptional network states underlying xenograft tumor cells recapitulate those states present in the primary tumor.

**Figure 2.**
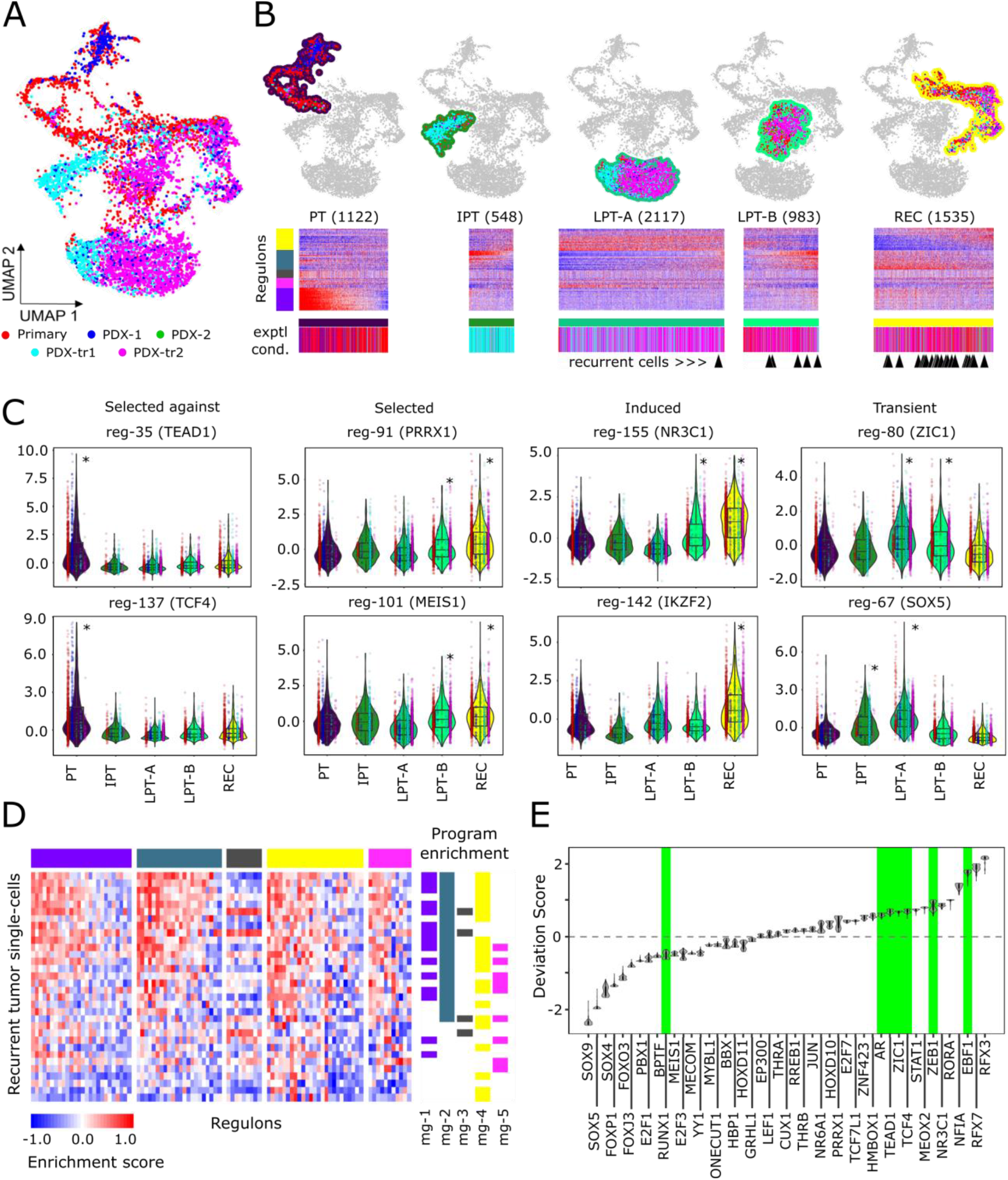
Modeling tumor progression and tumor response to standard of care. **(A)** UMAP plot of integrated scRNA-seq profiles of tumor cells from UW7 primary tumor and corresponding PDX mice models. Colors indicate experimental conditions (Primary tumor = tumor biopsy, PDX-1/2 = untreated samples collected at 24 hrs and 72 hrs post completion of corresponding TMZ/XRT treatment, PDX-tr1/tr2 = samples collected 24 hrs and 72hrs, respectively, post TMZ/XRT treatment). **(B)** Five subpopulations of tumor cells enriched for various experimental conditions (PT = pretreatment, IPT = immediate post-treatment, LPT-A = late post-treatment A, LPT-B = late post-treatment B, REC = recurrent). Heatmaps underneath each subpopulation show regulon activity z-scores across single-cell samples for each subpopulation. Color bars underneath each heatmap indicate the experimental condition of each cell as in A. Black arrowheads indicate which tumor cell sample the recurrent tumor cells collected at autopsy mapped closest to in the alternative UMAP embedding plot (Fig. S5). **(C)** Violin plots of regulon activity within each subpopulation. Select examples of regulons exhibiting distinct dynamic behavior, (i.e., “selected-against”, “selected”, “induced”, and “transient” behavior) are included. Inline points represent regulon activity levels within cells from each experimental condition within each subpopulation. Asterisks indicate which subpopulations had significantly higher regulon activity relative to the rest of the primary/PDX tumor-cell population (FDR-adjusted *p* value < 0.05). **(D)** Gene set enrichment analysis of regulon gene sets in recurrent tumor cell scRNA-seq profiles. Top color bar indicates transcriptional programs (Fig. 1D). Right adjacent color bars indicate samples statistically enriched with regulons for a particular transcriptional program (FDR-adjusted p-values ≤ 0.1). **(E)** Violin plots showing deviation scores across recurrent tumor scATAC-seq profiles. Green background highlights the 7 consensus TFs previously identified (Fig. 1H).

In addition, our framework enabled us to identify patterns of regulon/transcriptional program activities that were reflected in the recurrent tumor. We first compared scRNA-seq profiles of tumor cells identified from the recurrent tumor specimen collected at autopsy (Fig. S4) to those of the primary tumor and PDX samples. Specifically, we projected the recurrent tumor cells into a low dimensional space (Fig. S5) defined by the primary and PDX tumor cells and found that a majority of the recurrent tumor cells projected onto primary tumor and PDX cells in SNN cluster 1 (Fig. S5). Concomitantly, we used single sample gene set enrichment analysis (ssGSEA) to test for the enrichment of regulons and transcriptional programs within the autopsy tumor cells. Indeed, a majority of autopsy tumor cells were significantly enriched for transcriptional programs that were active in late-stage post-treatment PDX samples (Fig. 2D) and shared similar transcriptomic states and programmatic activity to those tumor cells as well (Fig. S6), which further support the utility of PDX models to characterize treatment-induced evolution of tumor cell states.

We examined the distribution of cells from different timepoints across the co-embedded UMAP space and identified four broad partitions in the data (Fig. 2B). The “early” (pre-treatment) stage was heavily enriched for cells from the primary tumor and untreated PDX models (Fig. S7). The “immediate post-treatment” stage (IPT) was enriched for the 24-hour post-treatment timepoint. The “late post-treatment” (LPT) stage was divided into two subpopulations (A and B) that were enriched for the 72-hour post treatment timepoint. Finally, the “recurrent” timepoint (REC) included cells from the recurrent, autopsy specimen and 72-hour post treatment PDX samples.

### scSYGNAL/Open Targets platform analysis allows for identification of possible drug targets for induced, selected, and recurrent cell states

Understanding tumor evolutionary trajectories could have important implications for therapeutic strategy. Therefore, we sought to identify potential therapeutics that putatively target TF regulators and/or their associated regulon gene members exhibiting positive activity across tumor cell states within specific timeframes (Table S12) using the Open Targets platform (*30*). Briefly, the Open Targets platform provides an extensive curated database that enables users to identify drug-target genes/proteins pairings. Using this platform, we identified a set of drugs that target various regulators and/or downstream target genes associated with the regulons/programs identified in our analysis (Table S13). In addition to this, we curated the literature to identify pathways and drugs that were relevant to TFs in each of the trajectories. We then aligned these drug mappings with specific patterns of regulator activity over the timespan of our disease-modeling framework.

Using this framework, we characterized dynamics over the time course of treatment and recurrence with the aim of identifying potential therapeutic vulnerabilities and optimal treatment timeframes (i.e., regulon and cell state changes). Several dynamic patterns of cell state and specific regulon activity occur during the process of tumor evolution and offer an additional dimension along which to define treatment-related state and potentially inform treatment strategies. The first evolutionary path of a regulon to consider is one that is active early in the disease but decreases in activity over time. Our modeling system allows us to identify regulators that are represented only the in early (pre-treatment) tumor specimen. Of the consensus TFs identified from integrated analysis, *TEAD1* and *TCF4* demonstrated decreased activity after SOC (Fig. 2C), suggesting that SOC effectively depleted and selected against those regulons.

Another treatment-related state observed was the “selected” state, characterized by behavior in which regulons increase activity over the course of treatment. Thus, tumor cells within the primary tumor expressing these regulons in the primary tumor may have a relative advantage during treatment, which is supported by the subsequent increased activity and representation in cells over time. One example of a “selected” regulator is the transcription factor *PRRX1*, which had relatively low activity in the early stages but increased activity over the course of the disease (Fig. 2C). *PRRX1* has previously been associated with regulation of mesenchymal gene expression programs in cancer via activation of TGF-ß signaling (*31*). This would suggest that small molecule inhibitors of the TGF-ß signaling pathway such as galunisertib should have effects on this particular regulon. Because this regulon’s activity increases over the course of treatment, our data and analysis would suggest that TGF-ß inhibitors (i.e., galunisertib) should be combined with XRT/TMZ concurrently or could serve as an effective adjuvant therapeutic after SOC treatment.

In contrast, “induced” states are characterized by regulons having low activity in the early stages of the disease but become prevalent over time. *IKZF2*, a chromatin remodeler that has been shown to regulate chromatin accessibility in leukemia, was less active in the primary tumor cells but became active during treatment and remained active at recurrence (Fig. 2C). Open Targets analysis identified CDK4/6 inhibitors as possible therapeutics targeting this regulon through activity of *IKZF2* on CDK4 (Table S12). Once again, given the importance of these states during treatment, our analysis would suggest that therapies targeting this regulon are more likely to be effective in the concurrent and adjuvant setting. Induced states could also be identified that were unique to later stages. A regulatory network driven by *NR3C1* was independently identified from scSYGNAL analysis as a regulator of the recurrent stage and had one of the highest deviation scores from analysis of the ATAC-seq data (Fig. 2E). This would suggest that a regulatory network governed by *NR3C1* expression was particularly important at the time of tumor recurrence. Open Targets analysis identified corticosteroids, including dexamethasone, as possible modulators of this regulon. The activity of this regulon at the time of recurrence may be a reflection of dexamethasone administration at later stages of the disease. Moreover, the post-treatment upregulation of NR3C2, suggests that agonists such as eplerenone and spironolactone targeting this mineralcorticoid receptor could potentially be used in an adjuvant setting following SOC. Interestingly, previous studies have shown spironolactone to have anticancer properties in prostrate and breast cancer via regulation of DNA damage response (*32*).

### scSYGNAL allows for identification of transient therapy-induced evolutionary cell states and associated regulons

Finally, a key strength of this modeling framework was the ability to identify regulons and cell states that are only transiently-induced during treatment but do not persist beyond the immediate treatment or late post-treatment stages. These cell states are presumably missed by most clinical specimen analysis because they exist transiently during a time-period when sampling rarely occurs. In our framework, transient states are represented most clearly by the immediate post-treatment (IPT) and late post-treatment (LPT) timepoints. We identified *ZIC1* as being predominantly active during this part of the overall tumor evolutionary trajectory (Fig. 2C). *ZIC1* is a zinc finger TF that has been shown to regulate forebrain development by maintaining neural precursor cells in an undifferentiated state (*33*). The regulon for *ZIC1* includes several genes important in glioma stemness including *ADM* and *VGF*. Its activation during treatment suggests that targeting this regulatory network in the adjuvant setting may effectively block a pathway to resistance.

Another set of cell states and associated regulons that showed transient induction were regulated by *SOX5* (Fig. 2C), which has been shown to have mixed roles in glioblastoma but is an important driver of TGF-ß mediated epithelial-to-mesenchymal transition and invasive phenotypes in a variety of other cancers (*34*–*36*). Several regulons driven by *SOX5* were activated throughout the treatment process, including regulons specific to “immediate post-treatment” and “late post-treatment” stages. In addition to the TGF-ß pathway, Open Targets analysis also identified *TTK*, a dual specificity protein kinase as a member of the *SOX5* regulon. While there are no TTK inhibitors currently approved, there are several in development for solid cancers (*37*). Transient states that are governed by TFs such as *ZIC1* and *SOX5* represent appealing targets for concurrent therapies that could be trialed in conjunction with SOC.

When assessing potential targets, a critical point to consider is that multiple regulons may be controlled by a single master regulator yet have activities that are contextually disparate. One such example involves the androgen receptor (*AR*), which putatively regulates three regulons (Table S2, Fig. S8). Regulon 15 demonstrated “selected against” behavior, suggesting that SOC was effective at suppressing its activity. Regulon 20 demonstrated “transient” behavior, suggesting that it may have some activity during SOC and may relate to resistance to radiation, which has been suggested previously (*38*). Finally, regulon 17 demonstrated “induced” behavior and had high activity at recurrence. Activity of the AR regulon raises the possibility of androgen targeting therapies as possible treatments for GBM in a variety of contexts, including in concurrent with SOC therapy or in the adjuvant and salvage settings. Determination of whether this is a gender-specific effect (this patient was male) and whether *AR* activity is generalizable across patients will require analysis of additional samples.

Furthermore, to increase the number of potential regulatory targets, it is also important to consider regulators that were identified uniquely by either transcriptional or epigenetic analysis. For example, analysis of scATAC-seq data from the recurrent sample identified several TFs with highly significant motif deviations, mostly notably *RFX3* and *RFX7* (Fig. 2E). These two TFs play roles in ciliogenesis in the central nervous system, which has been identified as an important pathway in regulating glioblastoma growth and resistance to therapy (*39*). Importantly, these two regulators were not identified in transcriptomic network analysis and would have otherwise been missed had our analysis utilized scRNA-seq alone, highlighting the importance of multi-omic information to identify both shared and unique regulatory networks in different contexts. While there are no small molecules that currently target these regulators, they may be appealing candidates for nonconventional or novel directed therapies including CRISPR-Cas9 or antisense RNA-based approaches in the recurrent or salvage therapy setting.

## Discussion

In summary, tumors are complex ecosystems of cell types and various tumor-cell subpopulations interacting with one another in a complex microenvironment, mandating the use of single cell methods to understand tumor biology more precisely. In this study, we characterized a patient’s tumor, analyzing the transcriptomic and epigenetic state of tumor cells and corresponding microenvironment. By integrating multimodal single-cell analysis and *a priori* knowledge of regulatory relationships, enabled by SYGNAL and MINER analysis, we identified regulon-based tumor-cell subpopulations and underlying regulatory relationships, inferred from scRNA-seq data and corroborated by scATAC-seq analysis. Our modeling/monitoring framework demonstrated the ability to capture spatiotemporal tumor heterogeneity and underlying mechanistic drivers within regulatory networks based upon both transcriptomic and epigenetic states. Furthermore, PDX modeling revealed multiple potential trajectories of tumor cell progression in response to chemoradiotherapy and in the setting of recurrence.

Our proof-of-concept work herein provides the basis for the development of a modeling and analytical system that enables single-cell characterization of an individual patient’s tumor and inferred therapeutic vulnerabilities (Fig 3). Although further validation is required, in the form of *in vivo* studies of these putative druggable targets, our preliminary analysis and results suggest that systems biology techniques can be used to infer and predict therapeutic vulnerabilities that are either selected or induced during SOC treatment. Ultimately, the information gathered from such systematic modeling and analysis of individual tumors may inform clinical treatment in a more targeted manner and enable a rational, tailored precision medicine that accounts for intratumoral cell heterogeneity.

**Figure 3.**
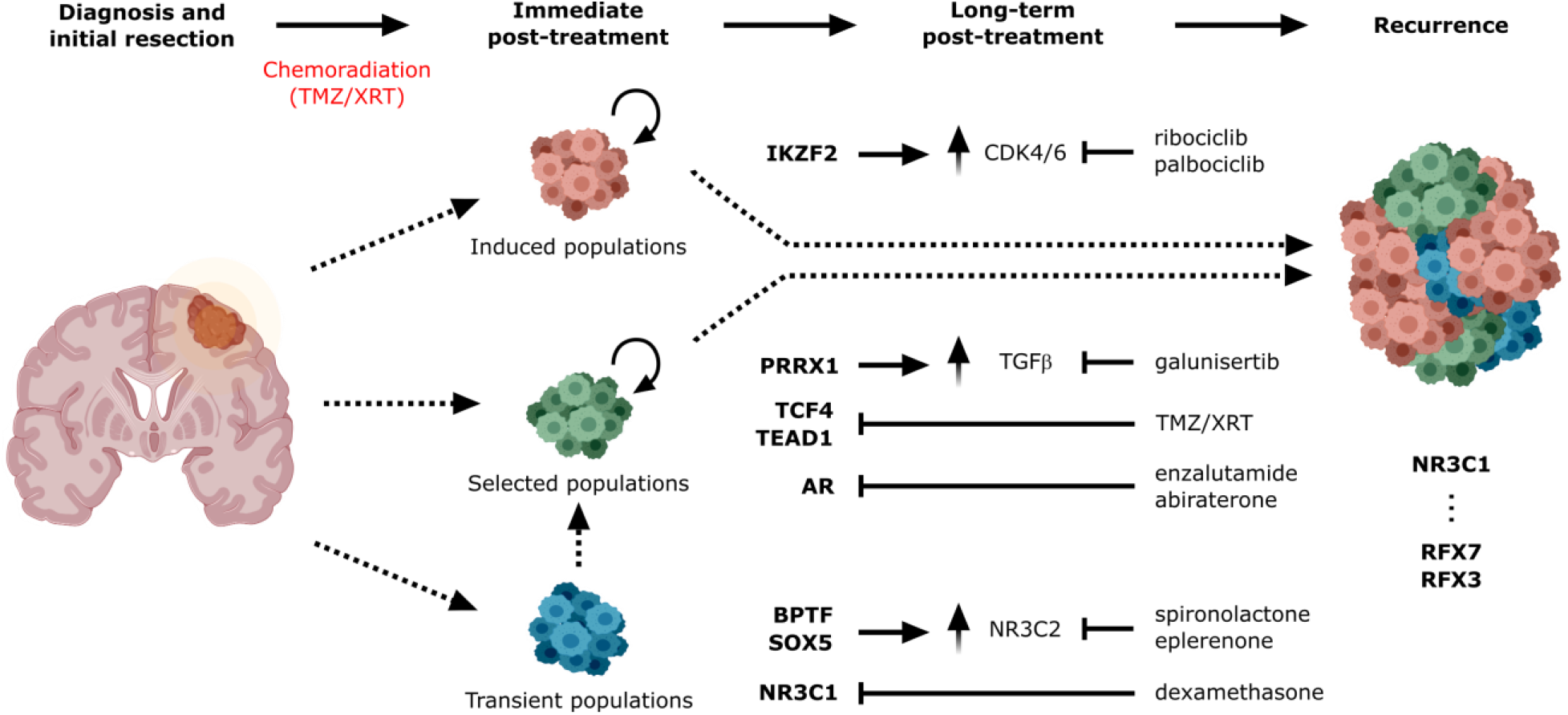
Comprehensive schematic of glioblastoma progression. Distinct populations of induced, selected, and transient tumor cells states, regulons, and TFs (bold) contribute to intratumoral heterogeneity that plays a role in treatment resistance. As cell states may be differentially susceptible to treatment and may be selected for or induced by therapeutic intervention, use of a more complete view of cell state trajectories with scSYGNAL and MINER analysis may allow for the prediction of therapies that work in either the concurrent setting against cell states or an adjuvant/neo-adjuvant setting against induced cell states.

## Materials & Methods

### Tumor acquisition

Based upon institutional review board (IRB)-approved protocols (protocol #STUDY00002162), intraoperative tumor specimens from adult patients who voluntarily consented to donation to the institutional tumor bank were collected in cryogenic vials (Corning; Corning, NY) and immediately snap frozen in liquid nitrogen. All patient specimens were anonymized prior to processing. Tumor pathology and diagnosis was confirmed by a neuropathologist as WHO grade IV glioblastoma, IDH-wild type. Specimen was then subsequently stored in −80 C freezers for further experimentation. Autopsy tissue was collected after informed consent with a waiver from the University of Washington IRB with a post-mortem interval of approximately 8.75 hours. Tissue was snap frozen in liquid-nitrogen cooled isopentane. Tumor regions were sampled based on gross examination of brain sections and processed as outlined below.

### Tissue processing

Frozen tissue was processed to nuclei using the Frankenstein protocol from Protocols.io. Briefly, snap frozen glioblastoma tissue was thawed on ice and minced sharply into <1 mm portions. 500 μl chilled Nuclei EZ Lysis Buffer (Millipore Sigma, NUC-101 #N3408) was added and tissue was homogenized 10-20 times in a Dounce homogenizer. The homogenate was transferred to a 1.5 ml Eppendorf tube and 1 mL chilled Nuclei EZ Lysis Buffer was added. The homogenate was mixed gently with a wide bore pipette and incubated for 5 minutes on ice. The homogenate was then filtered through a 70-μm mesh strainer and centrifuged at 500g for 5 minutes at 4°C. Supernatant was removed and nuclei were resuspended in 1.5 mL Nuclei EZ lysis buffer and incubated for 5 minutes on ice. Nuclei were centrifuged at 500g for 5 min at 4°C. After carefully removing the supernatant, nuclei were washed in wash buffer (1x PBS, 1.0% BSA, 0.2 U/μl RNase inhibitor). Nuclei were then centrifuged and resuspended in 1.4 ml wash buffer for two additional washes. Nuclei were then filtered through a 40 μm mesh strainer. Intact nuclei were counted after counterstaining with Trypan blue in a standard cell counter.

### Animal models

All animal procedures were performed in accordance with protocols approved by the Institutional Animal Care and Use Committee (IACUC) at Fred Hutchinson Cancer Center and the University of Washington. Animals were housed at a maximum of five per cage with 14-hour light/10-hour dark cycle with food and water *ad libitum*. Female 4-8 week-old NOD-SCID mice (NOD.Cg-Prkdcscid Il2rgtm1Wjl/SzJ, Jackson Labs; Bar Harbor, ME) were used for all experiments with random assignment into treatment groups where applicable. Mice were monitored at least three times weekly for weight loss and other signs of neurologic or physical distress.

### Patient-derived xenograft modeling

Fresh surgically resected tumor sample was placed in sterile phosphate buffered saline and transported to Fred Hutchinson Cancer Center for further processing. Tumor specimen was dissociated with the use of a papain-based tumor dissociation kit (Miltenyi Biotec, 130-095-942) as per manufacturer’s instructions. Intracranial orthotopic transplantation of single-cell suspension human glioblastoma tumor cells into murine mouse models were performed in standard, IACUC-approved fashion. Briefly, mice were induced with 5% isoflurane and maintained at 2% isoflurane in oxygen thereafter. After appropriate placement on a stereotactic frame (Stoelting Co.), the skull of the mouse was exposed through a small skin incision, and a small 1 mm^2^ burrhole was placed shortly behind and lateral to bregma using a 25-gauge needle. Freshly-dissociated cells were suspended in 5 mL of PBS and loaded into a 33-gauge Hamilton needle syringe. The cells were then subsequently injected 2.0 mm lateral and posterior to bregma and 2 mm deep to the cortical surface. After completion of injection, the syringe was left *in situ* for another minute before removal in attempt to minimize risk of cell reflux. After scalp closure with suture, the mice were removed from anesthesia and allowed to recover on warming pads and returned to their cages following full recovery. Mice were then checked daily for five consecutive days for signs of distress or neurologic disability. Mice were also monitored using a small animal 1.5T MRI to track the degree of intracranial tumor, initially four weeks following injection and then again upon signs of neurologic symptoms, including ataxia, head tilt, seizures, or cachexia. Mice were sacrificed as soon as they demonstrated symptoms, and their brains were collected directly following euthanasia.

### Radiation and TMZ treatment

Tumor-bearing mice, as confirmed by small animal MRI, were given 50 mg/kg of temozolomide dissolved in 5% DMSO/saline or vehicle intraperitoneally for five consecutive days. On the same days, tumor-bearing mice were sedated with ketamine and xylazine and irradiated using a X-RAD 320 from Precision X-Ray at 115 cGy/min as has been performed previously (*40*).

### 10x Chromium scRNA-seq & scATAC-seq

Single-cell RNA sequencing was performed using the 10X Chromium v2 system. Library preparation was performed using 10x manufacturer instructions on an Illumina NovaSeq 6000. scATAC-seq was performed as per manufacturer instructions (SingleCell_ATAC_ReagentKits_v1.1_UserGuide_RevD) and sequenced on an Illumina NovaSeq 6000.

### Cell hashing and demultiplexing

Single nuclei from each PDX condition were labeled with 1 μl condition-specific hashtag oligonucleotide-labeled antibodies (BioLegend, TotalSeq A0541-A0545) according to manufacturer’s protocol prior to pooling and loading on a single lane of the 10X Chromium v2 system. The HTO library was processed separately and spiked in at 10% of the mRNA library prior to sequencing. Demultiplexing of pooled single-cell samples relies on subsequent HTO raw counts generated from scRNA-seq to classify computationally single-cells in their appropriate experimental condition. Demultiplexing was performed using the HTODemux function in the Seurat v3 platform (*9*). The result is single-cell annotation indicating the experimental condition in addition to the potential doublet or untagged state to which each tagged (or untagged) cell belongs.

### Doublet prediction

For those cells not processed using cell hashing (i.e., UW7 parental and UW7 recurrent autopsy cells), an alternative, computationally-based approach known as DoubletDecon was used to identify likely doublet samples (*41*). Briefly, DoubletDecon generates synthetic doublets by merging transcriptional profiles from randomly-selected pairs of cells belonging to distinct clusters identified in the dataset. These synthetic doublets are used, in conjunction with the previously identified clusters to create a deconvolution cell profile for the entire cell population. Pearson correlations are then calculated between each DCP and the centroid of each cluster. Those cells having the highest correlation to clusters comprised of synthetic doublets are labeled as doublets. Prior to final labeling of cells, a rescue step is performed in which certain cells may be rescued from the doublet labeling if the cell contains statistically significant upregulated expression, relative to a synthetic doublet cluster, for a minimum number of genes, that those cells are reincorporated into the non-doublet population. Finally, due to the random nature of synthetic doublet, it is likely that doublet predictions will vary run-to-run. Therefore, we conducted 50 runs to identify a consensus set of predicted doublets, which were subsequently excluded from downstream analysis.

### Quality control and scRNA-seq data pre-processing

We initially processed the 10X Genomics raw data using Cell Ranger Single-Cell Software Suite (release 3.1.0) to perform alignment, filtering, barcode counting, and UMI counting. Reads were aligned to the GRCh38 reference genome using the pre-built annotation package download from the 10X Genomics website. We then aggregated the outputs from different lanes using ‘cellrange aggr’ function with default parameter settings.

Each sample set analyzed via scRNA-seq (UW7 parental tumor, UW7 PDX samples, and UW7 recurrent tumor collected at autopsy) was QC-filtered separately prior to data integration, as in the case of UW7 parental tumor and UW7 PDX samples, and/or subsequent downstream analysis. Each sample set consisted of the following: 5,082 cells with 27,763 mapped genes (UW7 parental tumor), 11,648 cells with 26,231 mapped genes (UW7 PDX samples), and 690 cells with 19,917 mapped genes (UW7 recurrent autopsy tumor). To minimize inclusion of poor-quality genes and single-cell samples per sample set, we applied the following QC filters: 1) mapped genes must be expressed at minimum count of 2 in at least 20 cells, 2) mitochondrial genes must comprise ≤ 20% of the number of uniquely mapped genes/cell, 3) total counts/cell should be ≥ 500 and ≤ 50,000 (UW7 primary tumor cells), ≥ 500 and ≤ 24,000 (UW7 PDX samples), or ≥ 500 and ≤ 4,000 (UW7 recurrent autopsy tumor), and 4) the total number of mapped genes should be ≥ 500 genes and ≤ 10,000 (UW7 primary tumor cells), ≥ 500 genes and ≤ 7,000 (UW7 PDX tumor cells), or ≥ 500 genes and ≤ 30,000 (UW7 recurrent autopsy tumor). Post QC-filtering, each sample set consisted of the following: 4,456 primary tumor cells expressing up to 19,228 genes, 4,388 PDX tumor cells expressing up to 26,231 genes, and 350 recurrent tumor cells expressing up to 12,463 genes.

### Data normalization of scRNA-seq data

We applied the SCTransform function, provided in the Seurat v3.2.2 platform, to normalize and variance-stabilize UMI counts in the single-cell data. This function develops a regularized negative binomial regression model to characterize the UMI count distribution on a gene-by-gene basis. This model is then used to determine Pearson residuals (i.e., the square-root of the variance-normalized difference between the actual gene count and model-predicted counts). These residuals represent the standardized expression values not affected by technical artifacts and are used for downstream analysis. Concomitantly, mitochondrial gene expression influence was regressed out of expression for each gene in each cell, as part of the SC-normalization procedure.

### Batch integration of scRNA-seq data

To integrate the two different scRNA-seq datasets, we utilized the suite of integration functions provided by Seurat v3 platform – *FindTransferAnchors* and *IntegrateData*. These functions apply canonical correlation analysis (CCA) to identify shared patterns in gene expression profiles between datasets, (i.e., “integration anchors” that are pairs of cells that share maximal correlation with one another). These anchors are then used as references with which the remaining datasets are harmonized with one another. This technique enables information to be transferred across datasets including both continuous and categorical data, Consequently, categorical data like cell-type annotation or cluster membership can be transferred to multiple datasets that will be integrated together. We apply this method to transfer cell-type annotation from scRNA-seq data to corresponding scATAC-seq data (Fig. 1C).

### scRNA-seq cell-type and tumor cell annotation

Established CNS cell-type-specific genes were used to determine gene set module scores for each cell. Gene module scores were determined using the *AddModuleScore* function provided in Seurat. In brief, the module score represents the difference between the mean expression of the gene set of interest and the average expression of a randomly selected set of control genes. To create a set of control genes, all genes are first grouped into 25 bins according to their respective average expression. Next, for each gene in the gene set of interest, a corresponding set of 100 randomly selected genes is selected from the same expression bin. This results in a control set that is 100-fold larger in size, which is analogous to averaging over 100 randomly-selected gene-sets identical in size to the gene set of interest. Positive module scores indicate that the gene set of interest has higher expression than what is expected by random chance and vice versa. Final cell-type assignment was based on which corresponding gene set resulted in the highest positive module score above a threshold value of 0.1. To annotate tumor cells, inferCNV was used to infer the copy number variation state of each cell (Supplemental Text). Both cell-type and tumor cell state, defined by Chr7 gain and Chr10 loss, were used to determine final cell-type annotation for the primary and recurrent tumor biopsy samples (Figs. S1, S4).

### scRNA-seq multivariate analysis

Downstream analysis of scRNA-seq data was performed using Seurat v3.2.2. Following QC filtering, SC-normalization and integration, we performed principal component analysis (PCA) on the integrated gene expression matrix using the first 30 principal components for clustering and visualization. Next, we used the transformed gene expression data along the top 30 principal components (PC scores) to identify shared nearest neighbors (SNN). We then identified clusters in an unsupervised clustering using Leidan clustering using a resolution of 0.8. Visualization was performed using uniform manifold and projection (UMAP) using the scores values from the top 30 principal components using a minimum distance of 0.2 and a spread value of 1.2.

### Quality control and scATAC-seq data preprocessing

Similar to scRNA-seq data, we initially processed 10X Genomics raw data using the Cell Ranger Software Suite (release 3.1.0). We performed additional data preprocessing and analysis using the software package ArchR (version 0.9.5). As part of the QC-filtering process, we used 2 metrics including: 1) number of unique nuclear fragments (>1000), and 2) signal-to-background ratio (i.e., transcription start site (TSS) enrichment score > 4). This score represents a ratio of per-basepair accessibility centered around the TSS relative to flanking regions (2000 bp distal in either direction). Here, we used a TSS enrichment score value of 4 as a lower limit threshold. We also inspected fragment size distribution to verify whether a periodicity in fragment size, reflected as a multimodal distribution, existed. These peaks and valleys in the distribution occur because fragments span 0, 1, 2, etc. nucleosomes and the Tn5 enzyme cannot cut DNA that is tightly wrapped around a nucleosome. Moreover, we inferred and removed likely doublets from the datasets. Doublet inference in ArchR involves a method similar to the DoubletDecon in that heterotypic doublets are synthesized from the original population. These synthetic doublets are then added to the original population, which is projected into a 2D space via UMAP (*42*). Single-cells are then labeled as putative doublets if they repeatedly project as nearest neighbors during this iterative procedure.

We calculated QC statistics separately for each scATAC-seq data set (UW7 parental tumor and UW7 autopsy single-cell samples). The UW7 parental tumor set initially included 3,770 cells, having a median of 31,462 fragments/cell. In this case, applying QC-filtering resulted in 3,407 cells having a median of 29,268 fragments/cell. The UW7 autopsy data set initially included 1,934 single-cells, with a median of 8,801 fragments/cell. Following QC-filtering, 1,425 single-cells with a median of 8,033 fragments/cell remained (Fig. S9, S10).

### scATAC-seq dimensionality reduction

Due to the sparse nature of scATAC-seq data, popular methods like PCA would result in high inter-cell similarity due to the predominance of non-values in the scATAC-seq profiles across the single-cell samples. Towards addressing the sparsity issue, latent semantic indexing (LSI), a technique applied in natural language processing to assess document similarity based on word counts, which often involves sparse and noisy datasets (many words, low frequency). Analogously, scATAC-seq profiles are viewed as a document and different accessible regions/peaks are words. To reduce the dimensionality of the scATAC-seq dataset, term frequency by depth normalization per cell is calculated. Next, these values are normalized by the inverse document frequency, which weights features by how often they occur. The result is a matrix that indicates how important a region/peak is to a sample. Using this resulting matrix, singular value decomposition (SVD) is applied to factorize the matrix into constituent matrices from which the most valuable information can be identified and projected into a lower dimensional space.

Here, ArchR applies a variation of this LSI methodology, an iterative LSI approach (*43, 44*). The default setting of two iterations was performed on both UW7 parental tumor and UW7 PDX scATAC-seq datasets.

### scATAC-seq cell-type and transcriptional program labeling

Labeling of scATAC-seq datasets was performed using ArchR (package 22, v0.9.4). In brief, filtered fragments.tsv.gz files after quality control were used to generate an ArchR GeneScore matrix and a tiled genome feature matrix for each dataset. Cells were grouped by performing iterative latent semantic indexing (LSI) on the tile matrix, followed by the shared nearest neighbor clustering approach implemented in Seurat v3.2.2. GeneScore data, a correlate for gene expression, was then used to compare scATAC-seq clusters to a labeled reference scRNA-seq dataset, the UW7 parental tumor single-cell samples, using ArchR’s implementation of the *FindTransferAnchors* method from Seurat. Cell-type and/or sample groups based on transcriptional network states with the highest score for each scATAC-seq cluster were used to annotate those cells for downstream analysis and display (Fig. 1C-1H).

### Motif deviation scores

TF motif deviations were predicted on a per cell bases, relative to an aggregate background of a subpopulation of cells via chromVAR, which was incorporated into the broader ArchR package. The enrichment of TF motifs can guide in the prediction of which regulatory factors are most active in a cell type of particular interest, such as tumor cells. Designed for predicting enrichment of TF activity on a per-cell basis from sparse chromatin accessibility data, chromVAR produces 2 outputs including 1) deviation: a TN5 insertion sequence bias-corrected measurement of how far the per-cell accessibility of a given motif deviates from the expected accessibility based on the average of all cells or samples, and 2) z-score: referred to as a “deviation score” for each bias-corrected deviation across all cells. The absolute value of the deviation score is correlated with the per-cell read depth. With more reads, there is higher confidence that the difference in per-cell accessibility of the given motif from the expectation is greater than would occur by chance.

### Regulatory network analysis

To infer regulons within single cells, we applied the SYGNAL (*11*) and MINER (*12*) workflow to the scRNA-seq data set resulting from the Batch Integration procedure described above. The MINER algorithm involves a suite of functions that enables the inference of causal mechanistic relationships linking genetic mutations to transcriptional regulation. Because our datasets did not include any extensive mutational profiling, we primarily focused on identifying regulons, based on co-expression clustering and enrichment of transcription factor binding motifs present in those co-expression clusters identified, and calculated the activity of these regulons in the single-cell samples. Regulon activity represents the eigengene value in each individual cell. Briefly, regulons identified in part by performing PCA on the normalized scRNA-seq data profiles in a gene-centric manner, i.e., PCA is used to identify principal components in which decreasing amounts of variation *across genes* is captured along each principal component – defined as a linear combination of samples in this approach. Here the coefficients, i.e., loadings, associated with each sample making up a principal component represent the eigengene value (*45*). Alternatively, one can view eigengene values as a scalar representation of expression of gene members for a regulon.

To determine the significance of each inferred regulon, we performed a permutation test to determine the possibility of obtaining an eigenvalue corresponding to the 1st principal component of a regulon (across all single-cells) of equal or greater value. The eigenvalue represents a summarizing value of all the genes in the regulon, i.e., eigengene and thus if these genes are indeed share coregulation or are correlated, the eigengene value would be higher than that of randomly selected set of genes. Next, we randomly select a set of genes having the same number of members as the original regulon and calculate the corresponding eigengene value for the permuted regulon. This procedure is repeated 1000 times to create a null distribution of eigengene values. We repeated this procedure for each inferred regulon. Those regulons whose eigengene values were greater than the 95th percentile of their respective null distribution were considered significant. Furthermore, we used eigenvalues to represent regulon “activity” within each cell.

Using the calculated activities of regulons, we identified groups of regulons sharing similar activity profiles across the cell population, i.e., transcriptional programs. Specifically, those regulons that correlated across the cell population (k-means clustering of sample pairwise Pearson correlations) defined distinct transcriptional programs. We further defined subpopulations of single-cells based on their shared regulon/transcriptional program activity. Sample pair-wise Pearson correlations were calculated based on their regulon activity profiles.

### Regulon enrichment analysis

We used the geneset variance analysis GSVA (version 1.34.0, R package) (*46*) to determine enrichment scores of genesets. To confirm the significance of these enrichment scores, we performed permutation tests in which gene rankings were randomized in each single-cell sample and calculated the corresponding enrichment scores. In total, 1000 permutations were performed, from which the resulting scores were used to define empirically a null distribution of enrichment scores. We considered regulons having enrichment scores greater than the 95th percentile of the null distribution to be enriched in a particular cell.

### Projection of UW7 recurrent tumor cells onto UW7 primary and PDX tumor cell UMAP embeddings

Before projecting any new data onto pre-existing data, we first determined a common set of gene features across the datasets, from which dimensionality reduction and data projection could be performed. We identified the common 6,541 genes across all the datasets. We then repeated PCA on the integrated UW7 data set using only the 6,541 common set of genes and used the transformed gene expression data along the top 30 principal components for visualization via UMAP. We then mean-centered and variance-normalized the UW7 autopsy tumor cell expression data using gene-specific means and variances calculated from the integrated dataset. These mean-centered and variance-normalized values were transformed via matrix multiplication with the eigenvectors from the top 30 principal components. We used the *predict* function in R along with the UMAP embeddings for the integrated data set to develop a linear regression model and the UW7 autopsy transformed data as predictors. Once the UW7 recurrent tumor sample UMAP embeddings were determined, we calculated pairwise Euclidean distances in the UMAP space amongst all tumor cell pairs between the UW7 autopsy and integrated datasets. Those cells having the lowest distance to the projected UW7 autopsy tumor cells are represented as arrowheads in Fig 2B.

### Drug Matching Identification

To identify drugs targeting elements within the transcriptional programs and states identified from the network analysis, we applied the Open Targets platform tool (https://www.targetvalidation.org/). The platform integrates a variety of data and evidence from genetics, genomics, transcriptomics, drug, animal models, and literature to score and rank target-disease associations for drug target identification. We focused our search on identifying drug-target matches for only those drugs associated with any cancer treatment employed in clinical trials stages phase I through IV.

## Acknowledgments

We thank the University of Washington Brain Tumor Bank for collection and acquisition of freshly resected tumor samples as well as the patient UW7 and family for their generosity.

## Author contributions

S.N.E., A.M., A.H.F., and A.P.P. conceived the project and designed experiments. S.E. and A.M. developed the PDX mouse modeling system. A.P.P., C.D.K., and P.J.K. provided and curated the patient-derived primary, biopsy, and autopsy tumor specimen. A.H.F., S.N.E., and A.B.M. performed *in vitro* and *in vivo* experiments. J.P., A.L., K.K., W.W., S.T., N.S.B., and A.P.P. performed and/or contributed to data analysis. J.P., A.H.F., N.S.B., and A.P.P. drafted and edited the manuscript with input from all authors.

## Data and materials availability

Data to be submitted in GEO.

## List of Supplementary Materials

**Figure S1**. Tumor cell annotation in primary tumor biopsy.

**Figure S2**. Enrichment of experimental conditions within shared-nearest-neighbors (SNN) clusters of primary and PDX tumor cells.

**Figure S3**. Primary tumor cells and PDX tumor cell nearest neighbors.

**Figure S4**. Tumor cell annotation of recurrent tumor biopsy.

**Figure S5**. Projection of UW7 recurrent autopsy tumor-cell samples into lower dimensional space defined by primary and PDX tumor cells.

**Figure S6**. Cosine similarity of recurrent tumor cells to LPT-B and recurrent cell groups with respect to transcriptional program 2 activity.

**Figure S7**. Enrichment of experimental conditions within timepoint partitioned clusters.

**Figure S8**. AR-regulated regulon activity across partitioned clusters.

**Figure S9**. Quality control & doublet identification for scATAC-seq profiles from primary tumor biopsy.

**Figure S10**. Quality control & doublet identification for scATAC-seq profiles from recurrent tumor biopsy.

**Table S1**. Cell type marker differential expression analysis. See attached.

**Table S2**. Regulons. See attached.

**Table S3**. FIRM analysis/miRNAs. See attached.

**Table S4**. Functional enrichment in transcriptional program 1. See attached.

**Table S5**. Functional enrichment in transcriptional program 2. See attached.

**Table S6**. Functional enrichment in transcriptional program 3. See attached.

**Table S7**. Functional enrichment in transcriptional program 4. See attached.

**Table S8**. Functional enrichment in transcriptional program 5. See attached.

**Table S9**. Breakdown and enrichment of proliferating cells in sample groups. See attached.

**Table S10**. Comparison of regulon TFs and CRISPR-Cas9 knockout screen of 1,445 TFs that overlap with GBM SYGNAL network. See attached.

**Table S11**. Differential expression analysis of ArchR-MINER TFs. See attached.

**Table S12**. Open Targets results. See attached.

**Table S13**. Significantly upregulated activity of regulons in timeframe groupings. See attached.

