## Supplementary text for "A single-cell based precision medicine approach using glioblastoma patient-specific models"

**This PDF file includes:**

Supplementary Text
Supplemental figures and legends S1 to S10

### Supplementary Text.

#### *Identification of tumor cells based on inferred CNV state and gene marker expression.*

Copy number variation (CNV) states were inferred via inferCNV. Briefly, scRNA-seq profiles are analyzed for evidence of large-scale chromosomal CNV including gains or deletions of large segments of a chromosome or entire chromosomes altogether. Expression intensity of genes across positions of the genome are compared to those in a reference set of cells (i.e., neurons identified based on expression levels of neuronal gene markers *RBFOX3* and *MAP2*). CNV states were estimated by sorting genes by their chromosomal location. A moving average was subsequently calculated along each chromosome using a sliding window of 100 genes. Relative gene expression was capped to a minimum and maximum of [-3, 3] to minimize the impact of any particular gene. Because GBM tumor cells exhibit a characteristic gain in Chr7 and loss in Chr10, we compared the CNV state of Chr7 and Chr10 genes across all cells relative to the previously-identified reference set of neurons within the dataset. Specifically, we calculated the sum of CNV states for

Chr7 and Chr10 within each cell. We then defined a 95% confidence interval around the sample mean of CNV sums for Chr7 and Chr10 for the reference cell set using a student's t-test distribution ($\nu = 197 - 1$ degrees of freedom, based on the number of reference neurons identified). Those cells simultaneously having a sum of CNV states greater than the 95% confidence interval range for Chr7 and less than the 95% confidence interval range for Chr10 were tentatively annotated as GBM tumor cells.

These preliminary inferred CNV-based tumor cell assignments were then mapped onto processed scRNA-seq profiles embedded into a 2D UMAP space. We then compared this preliminary tumor cell annotation against additional metrics including: 1) expression levels for gene markers (*EGFR*, *SOX9*, and *PTPRZ1*) associated with GBM tumors and 2) relative proximity of tumor and non-tumor cells to one another across UMAP embeddings (i.e., cluster membership of cells). Using a combination of these quantitative metrics, we refined the preliminary cell annotation to define any clusters of putative tumor cells.

***Projection of UW7 recurrent autopsy samples onto UW7 primary and PDX tumor cells.***

To project new data onto the pre-existing UMAP space, a common set of gene features was required. Comparison of the final gene sets that passed QC filtering across the UW7 primary tumor, UW7 PDX samples, and UW7 recurrent tumor biopsy revealed that 6,541 genes were common across the three datasets. Having identified a common set of gene features, we then reprocessed the integrated UW7 parental- and PDX-tumor cells following a workflow similar to described in the Methods section. First, we performed PCA on the refined integrated dataset. Secondly, we performed dimensionality reduction on the rotated dataset (i.e., scores along the first 30 PCs via UMAP which embedded the scores into a lower 2D space).

Following the embedding of the UW7 primary and PDX tumor cells defined by the refined set of common gene features, we then embedded the UW7 recurrent tumor cells in the UMAP space. We first projected recurrent tumor cells into the PC space defined by the UW7 primary and PDX tumor cells so that the appropriate rotations (scores) of the recurrent tumor cells were used for UMAP embedding. We performed a gene-centric mean-centering and scaling of the normalized scRNA-seq profiles of the UW7 recurrent tumor cells using the means and standard deviations defined by the UW7 parental/PDX tumor cells. A recurrent tumor cell was projected onto the pre-existing PCA space, defined by the UW7 primary/PDX tumor cells, by calculating the dot product between its mean-centered and scaled gene expression profile and the eigenvector defining a particular principal component. The resulting scalar product represented the score value of that recurrent tumor cell along that particular principal component. These score values for the projected UW7 recurrent tumor cells along the first 30 PCs were embedded into the UMAP space via linear regression.

***ArchR quality control metrics of scATAC-seq data.***

We followed the guidelines outlined in the ArchR package to assess the quality of the scATAC-seq data obtained from UW7 parental and recurrent tumor biopsies. Per the ArchR platform, three metrics are used to assess quality of scATAC-seq samples: 1) number of unique fragments (not mapping to mitochondrial DNA) per cell, 2) signal-to-background ratio (i.e., transcription start site [TSS] enrichment score, and 3) fragment size distribution based on nucleosomal periodicity. Below we briefly summarize the reasoning behind each QC metric, which is described in greater detail by Granja *et al*. (*18*).

Number of unique fragments as QC metric enables the identification of cells that have a low number of unique fragments and will not provide enough data to make any meaningful assessment of chromatin accessibility states and are excluded from downstream analysis. TSS enrichment score is based on the idea that ATAC-seq data is universally enriched at gene TSS regions as compared to other genomic regions due to large protein complexes that bind to promoters. By looking at per-bp accessibility centered at these TSS regions, we see a local enrichment relative to the surrounding regions (1900-2000 bp distal in both directions). The ratio between the peak of this enrichment (centered at the TSS) relative to these surrounding regions represents the TSS enrichment score.

Fragment size distribution provides yet another method to assess quality, providing a way to perform a sanity-check on data quality. DNA wraps around nucleosomes in a patterned way, where ~147 bp of DNA wrap tightly around a nucleosome. These tightly wrapped DNA regions cannot be cut by the Tn5 transposase. Consequently, fragment size distributions will often be depleted that are of the length of DNA wrapped around a nucleosome. Moreover, a result of this patterned DNA wrapping around a nucleosome is a periodicity in the distribution of fragment sizes in the data. Thus, hills and valleys are expected to appear in the size distribution because fragments must span 0, 1, 2, etc. nucleosomes.

### Supplemental Figure and Legends

**Figure S1.**

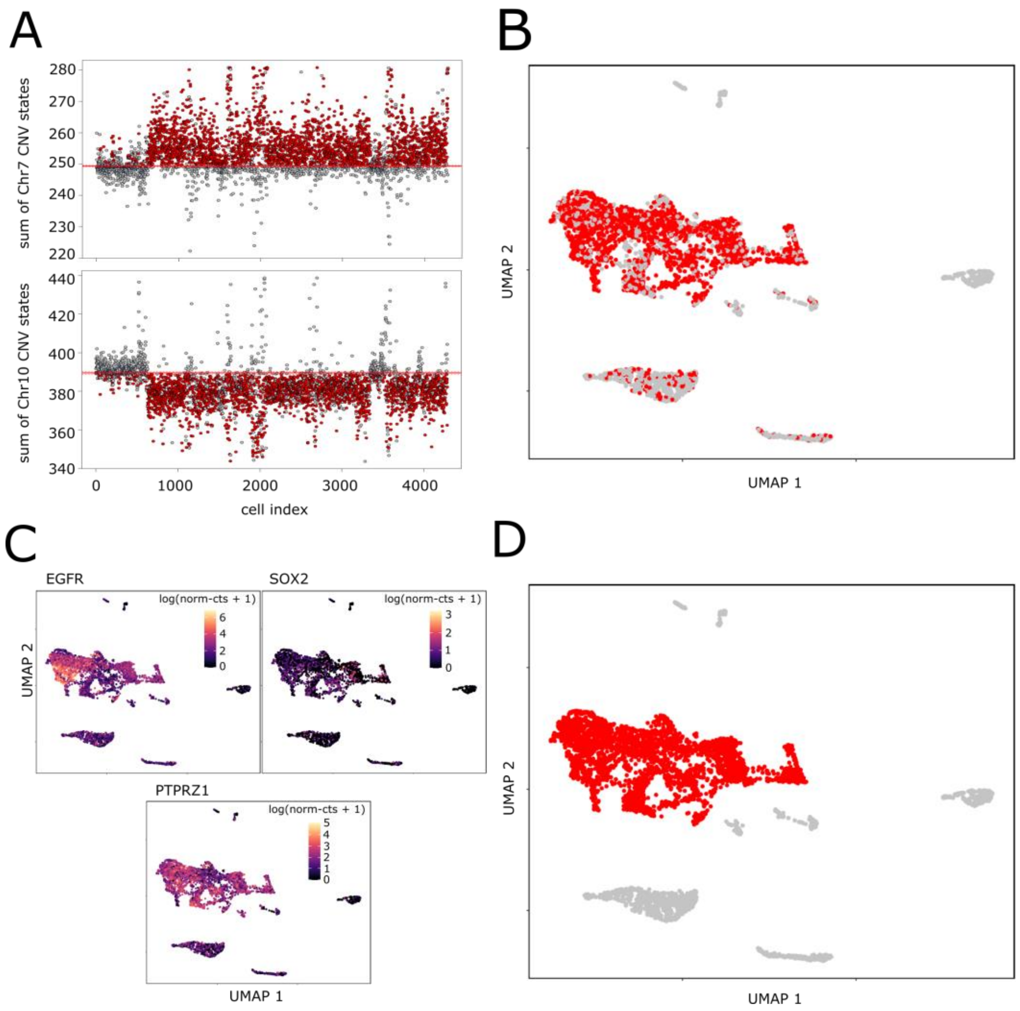


**Figure S1. Tumor cell annotation in primary tumor biopsy.** (A) Sum of inferred copy number variation (CNV) states for all cells for Chr7 and Chr10. Red solid lines represent the mean sum of inferred CNV state while dashed lines represent a 95% confidence interval of the average sum of inferred CNV state for reference neurons, identified based on their expression of neuronal markers RBFOX3 and MAP2. Red points represent cells having a Chr7 and Chr10 CNV state above and below the confidence intervals, respectively, based on CNV states of the reference cells. (B) UMAP visualization of UW7 primary tumor biopsy scRNA-seq profiles. Red colored cells represent cells having an inferred CNV gain in Chr7 and loss in Chr10, shown in (A). (C) UMAP visualization of UW7 primary tumor biopsy scRNA-seq profiles annotated according to log-normalized gene counts of tumor gene markers EGFR, SOX2, and PTPRZ1. (E) Final tumor-cell annotation of UW7 primary tumor biopsy. Based on the expression of tumor gene markers and CNV state, 3,130 cells of the total 4,183 cells analyzed from the parental tumor biopsy were defined as tumor cells.

**Figure S2.**

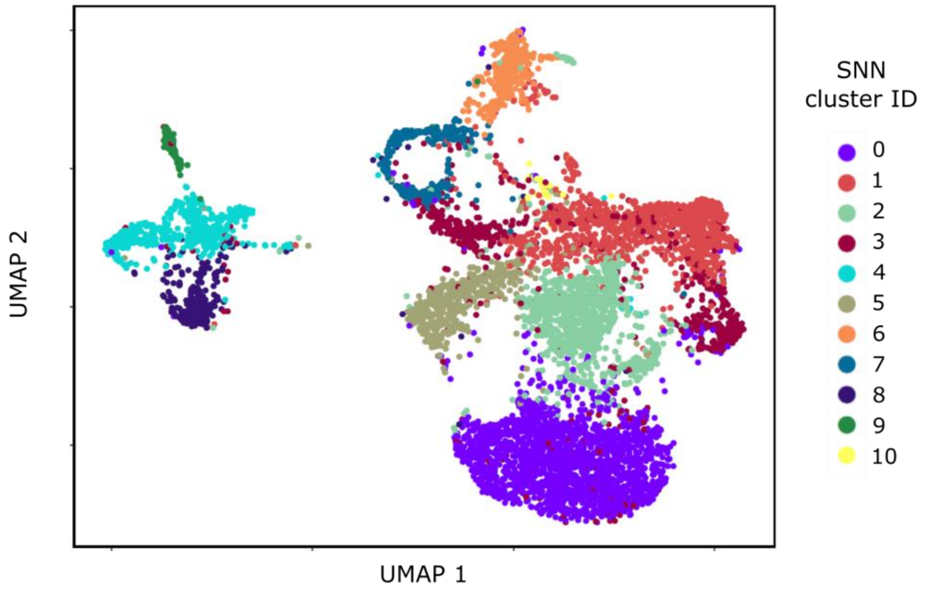


Breakdown of single-cell experimental conditions within SNN clusters

| | 0 | 1 | 2 | 3 | 4 | 5 | 6 | 7 | 8 | 9 | 10 |
|---|---|---|---|---|---|---|---|---|---|---|---|
| UW7 primary | 175 | 688 | 412 | 354 | 516 | 60 | 206 | 272 | 298 | 106 | 43 |
| UW7 PDX-1 | 72 | 39 | 20 | 18 | 2 | 20 | 184 | 21 | 0 | 3 | 2 |
| UW7 PDX-2 | 124 | 50 | 33 | 35 | 0 | 34 | 2 | 6 | 0 | 1 | 0 |
| UW7 PDX-tr1 | 427 | 107 | 47 | 64 | 2 | 411 | 0 | 14 | 4 | 3 | 2 |
| UW7 PDX-tr2 | 1354 | 312 | 442 | 232 | 33 | 10 | 0 | 12 | 0 | 5 | 0 |

Enrichement of single-cell experimental conditions within SNN clusters

| | 0 | 1 | 2 | 3 | 4 | 5 | 6 | 7 | 8 | 9 | 10 |
|---|---|---|---|---|---|---|---|---|---|---|---|
| UW7 primary | 1 | 1.45E-27 | 1 | 9.52E-05 | 5.32E-150 | 1 | 2.33E-4 | 5.55E-53 | 6.54E-105 | 4.59E-26 | 1.53E-11 |
| UW7 PDX-1 | 1 | 1 | 1 | 1 | 1 | 1 | 6.76E-144 | 0.598 | 1 | 1 | 1 |
| UW7 PDX-2 | 1.14E-06 | 1 | 1 | 0.276 | 1 | 1.17E-2 | 1 | 1 | 1 | 1 | 1 |
| UW7 PDX-T1 | 1.20E-13 | 1 | 1 | 1 | 1 | 7.02E-255 | 1 | 1 | 1 | 1 | 1 |
| UW7 PDX-T2 | 9.43E-262 | 1 | 1.43E-19 | 1 | 1 | 1 | 1 | 1 | 1 | 1 | 1 |

**Figure S2. Enrichment of experimental conditions within shared-nearest-neighbors (SNN) clusters of primary and PDX tumor cells.** Single-cell clusters identified using SNN modularity optimization and Leiden algorithm using the Seurat v3.2.2 platform. Top table tabulates composition of experimental conditions within each cluster. Bottom table tabulates FDR-adjusted p values for enrichment of experimental conditions within each cluster, per hypergeometric test.

**Figure S3.**

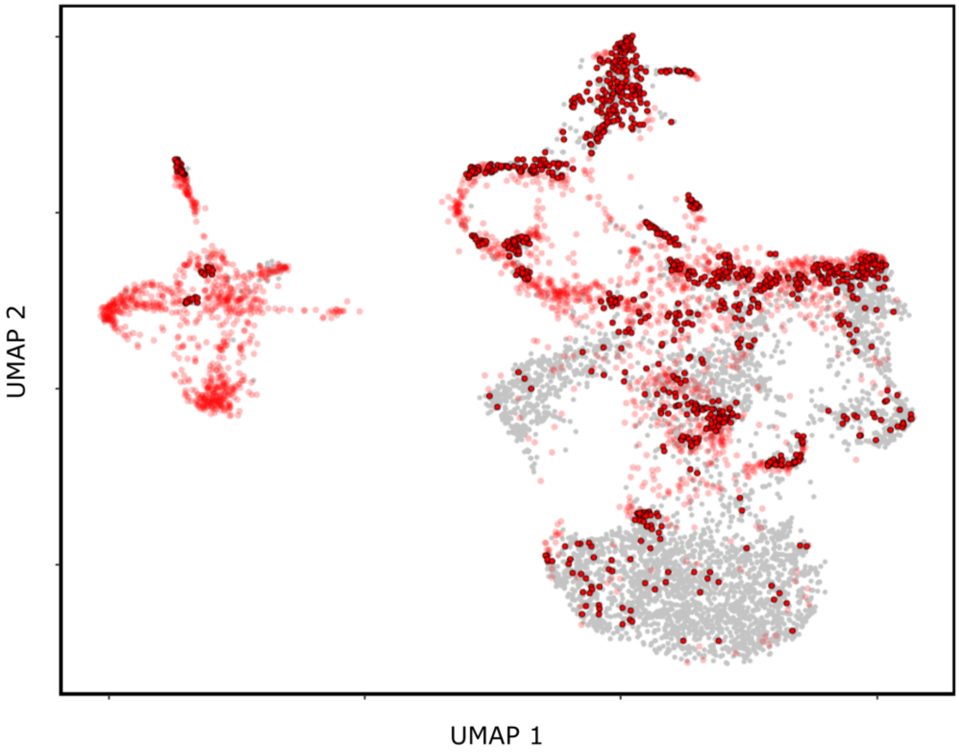


Breakdown of regulon-based transcriptional-network states across single-cells having PDX-sourced cell as nearest-neighbor

| Primary tumor cell category | Data-type | SG-1 | SG-2 | SG-3 | SG-4 | SG-5 |
|---|---|---|---|---|---|---|
| Primary tumor cells with PDX nearest-neighbor | Counts | 311 | 65 | 216 | 247 | 236 |
| | % of cells | 43.7 | 21.1 | 26.6 | 59.4 | 26.8 |
| All primary tumor cells | Counts | 712 | 307 | 813 | 416 | 882 |
| | % of cells | 22.7 | 9.8 | 26.0 | 13.3 | 28.2 |

**Figure S3. Primary tumor cells and PDX tumor cell nearest neighbors.** UW7 primary cells are highlighted in the UMAP visualization of UW7 primary and PDX single-cell samples. All red points represent UW7 primary tumor cells. Red points with black borders indicate which UW7 primary tumor cells have a PDX tumor cell within a radial distance – defined to be 1% of the maximal pairwise Euclidean distance amongst the UMAP embeddings. Table tabulates the composition of sample groups in which UW7 primary tumor cells having a PDX nearest neighbor belong.

**Figure S4.**

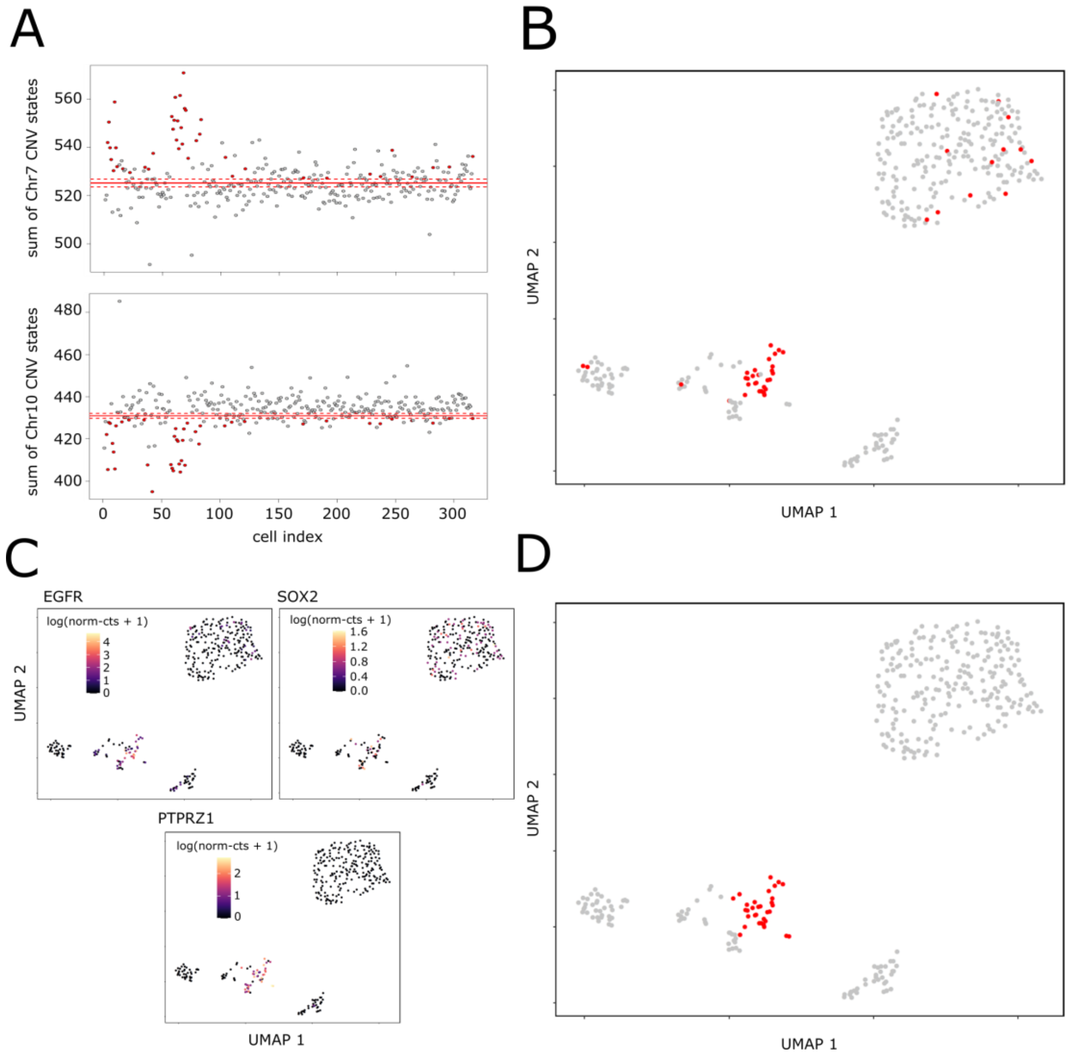


**Figure S4. Tumor cell annotation of recurrent tumor biopsy.** (A) Sum of inferred copy number variation (CNV) states for all cells for Chr7 and Chr10. The upper and lower red lines represent a 95% confidence interval of the average sum of inferred CNV state for reference neurons, identified based on their expression of neuronal markers RBFOX3 and MAP2. (B) UMAP visualization of UW7 recurrent tumor biopsy scRNA-seq profiles, annotated based on the preliminary tumor cell state defined in A. Red cells represent cells having an inferred CNV gain in Chr7 and loss in Chr10. (C) UMAP visualization of UW7 recurrent tumor biopsy scRNA-seq profiles annotated according to log-normalized gene counts of tumor gene markers EGFR, SOX2, and PTPRZ1. (D) Final tumor-cell annotation of UW7 recurrent tumor biopsy. Based on the expression of tumor gene markers and CNV state, 32 cells of the total 350 cells analyzed from the recurrent tumor biopsy were defined as tumor cells.

**Figure S5.**

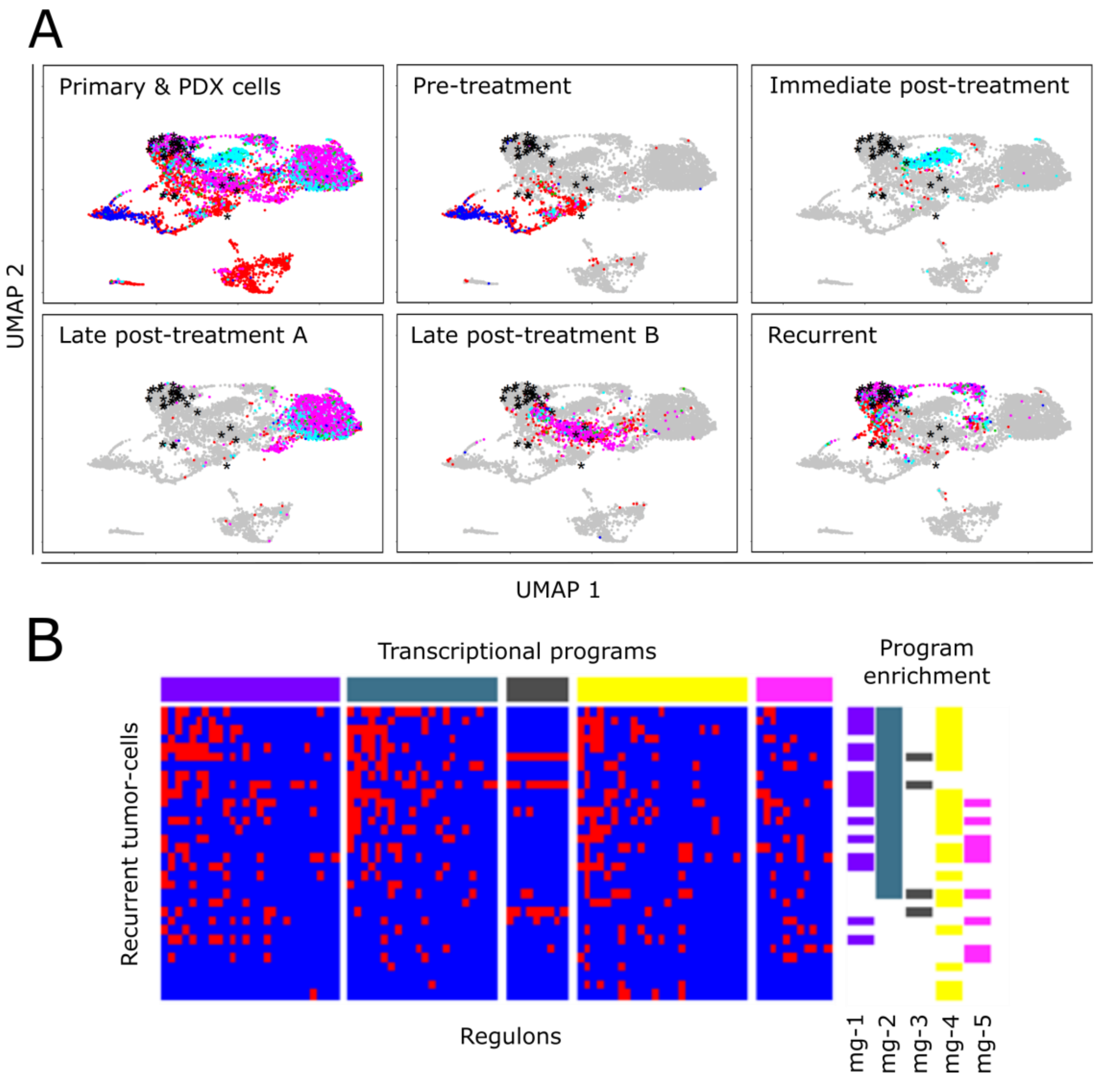


**Figure S5. Projection of UW7 recurrent autopsy tumor-cell samples into lower dimensional space defined by primary and PDX tumor cells.** (A) UMAP visualization of UW7 primary and PDX single tumor cells based on refined gene set common across UW7 primary/PDX single tumor cells and UW7 recurrent tumor cells collected at autopsy. Black astericks represent projected UW7 recurrent tumor cells. Color annotation for UW7 parental/PDX tumor cells is identical to that of Figure 2. Subpanels highlight those primary and/or PDX tumor cells belonging to the various timeframe groupings (Fig. 2B). (B) Heatmap visualizing p values for associated enrichment scores of regulons in UW7 recurrent tumor cells. Here, p values $\le$ 0.1 are highlighted in red while p values > 0.1 are in blue. Left-adjacent color bars mark recurrent tumor cells that exhibit statistical enrichment of regulons associated with a particular transcription program (Fig 1E), as determined by a hypergeometric test for enrichment of regulons having a positive and significant enrichment score.

**Figure S6.**

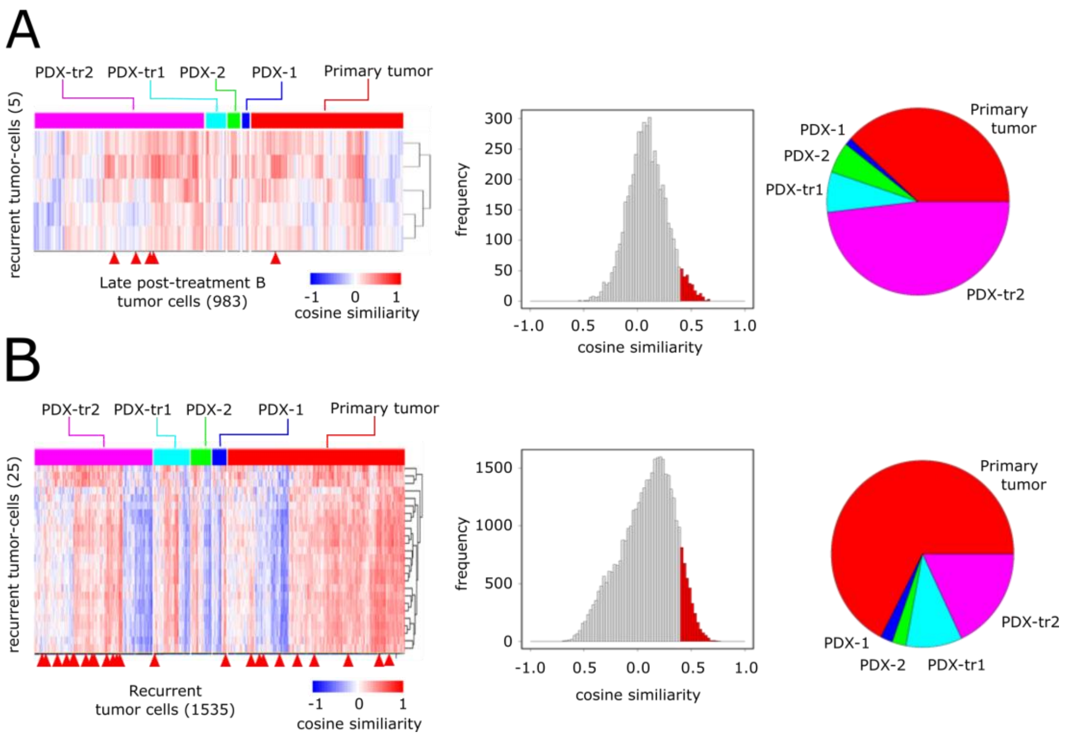


**Figure S6. Cosine similarity of recurrent tumor cells to LPT-B and Recurrent cell groups with respect to transcriptional program 2 activity.** (A) Heatmap of cosine similarity of UW7 recurrent tumor samples that projected onto UW7 primary/PDX tumor cells within the late post-treatment B timeframe (Fig. 2B). Red arrowheads underneath the heatmap mark those UW7 primary/PDX tumor cells that were the closest neighbors to the projected UW7 recurrent tumor samples in the UMAP embedding space. Histogram visualizes the distribution of cosine similarity scores (based on per-cell transcriptional program 2 activity, which is highly active in a majority of tumor cells within the 'recurrent' group – Fig 2B). The highlighted region indicate scores in the upper 90% quantile of the distribution. Adjacent pie graph indicates the breakdown of experimental conditions of UW7 primary/PDX cells with which the recurrent tumor cells share cosine similarity scores within the upper 90% quantile. (B) Similar series of graphs visualizing cosine similarity scores between UW7 recurrent tumor samples that projected onto UW7 primary/PDX samples within the recurrent timeframe (Fig. 2B).

**Figure S7.**

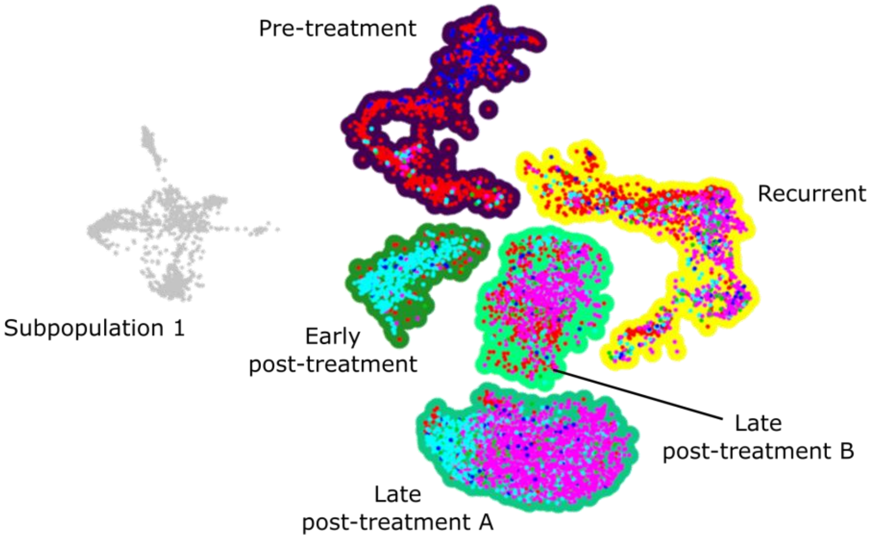


Breakdown of single-cell experimental conditions within timeframe groupings

| | Pre-treatment | Early post-treatment | Late post-treatment A | Late post-treatment B | Recurrent | Sub-population 1 |
|---|---|---|---|---|---|---|
| UW7_primary | 839 | 55 | 414 | 151 | 744 | 927 |
| UW7PDX-1 | 214 | 19 | 19 | 66 | 59 | 4 |
| UW7PDX-2 | 13 | 35 | 34 | 117 | 85 | 1 |
| UW7PDX-tr1 | 26 | 424 | 54 | 418 | 151 | 8 |
| UW7PDX-tr2 | 30 | 15 | 462 | 1365 | 496 | 32 |

Enrichment of single-cell experimental conditions within timeframe groupings

| | Pre-treatment | Early post-treatment | Late post-treatment A | Late post-treatment B | Recurrent | Sub-population 1 |
|---|---|---|---|---|---|---|
| UW7_primary | 3.95E-121 | 1 | 1 | 1 | 2.71E-06 | 0.00E+00 |
| UW7PDX-1 | 8.76E-80 | 1 | 1 | 1 | 1 | 1 |
| UW7PDX-2 | 1 | 0.006981818 | 1 | 2.37E-05 | 0.000753 | 1 |
| UW7PDX-tr1 | 1 | 6.19E-267 | 1 | 5.49E-13 | 1 | 1 |
| UW7PDX-tr2 | 1 | 1 | 2.98E-22 | 1.62E-283 | 1 | 1 |

**Figure S7. Enrichment of experimental conditions within timepoint partitioned clusters.** UMAP plot of single tumor cells and corresponding clusters identified based on experimental timepoints. Top table tabulates composition of experimental conditions within each cluster. Bottom table tabulates FDR-adjusted p values for enrichment of experimental conditions within each cluster, per hypergeometric test. Subpopulation 1 is a smaller population of tumor cell, predominantly comprised of UW7 primary tumor cells.

**Figure S8.**

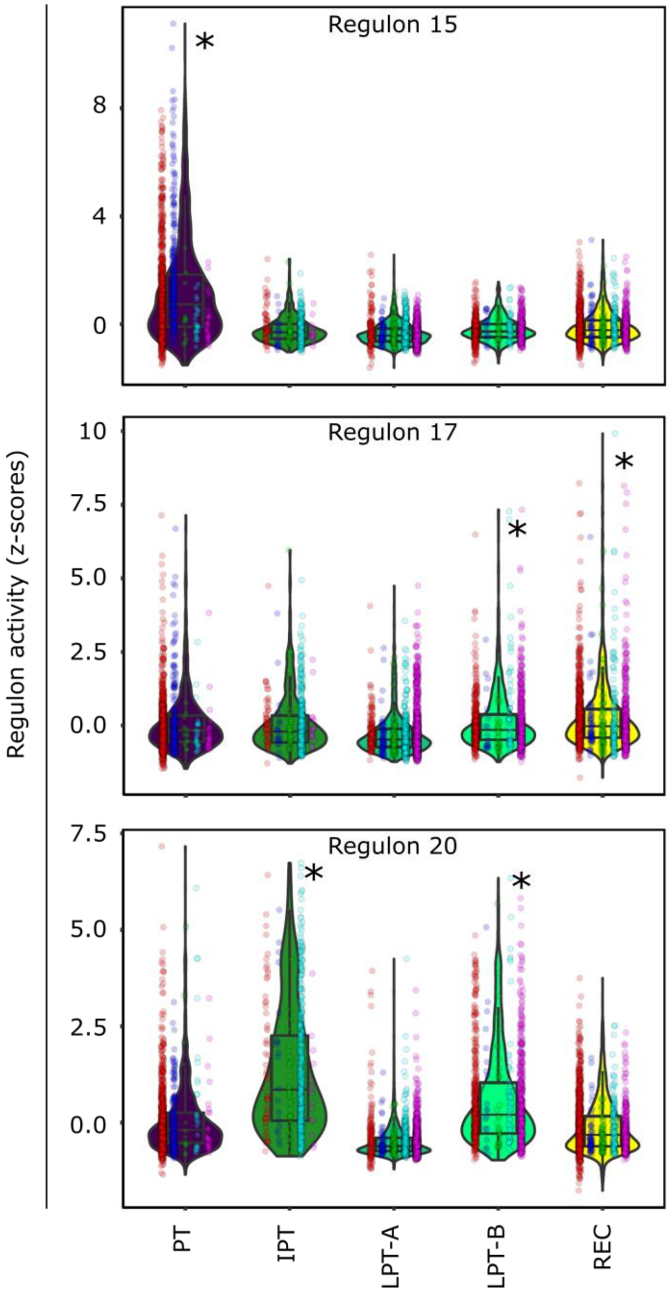


**Figure S8. AR-regulated regulon activity across partitioned clusters.** Violin plots of regulon activity within each subpopulation for AR-regulated regulons 15, 17, and 20. Regulon 15 exhibited "selected against' behavior across the time course of treatment. Regulon 17 exhibited 'induced' behavior and regulon 20 exhibited transient behavior. Asterisks indicate which subpopulations had significantly higher regulon activity relative to the rest of the primary/PDX tumor-cell population per Wilcoxon rank sum test (FDR-adjusted p value << 0.05).

**Figure S9.**

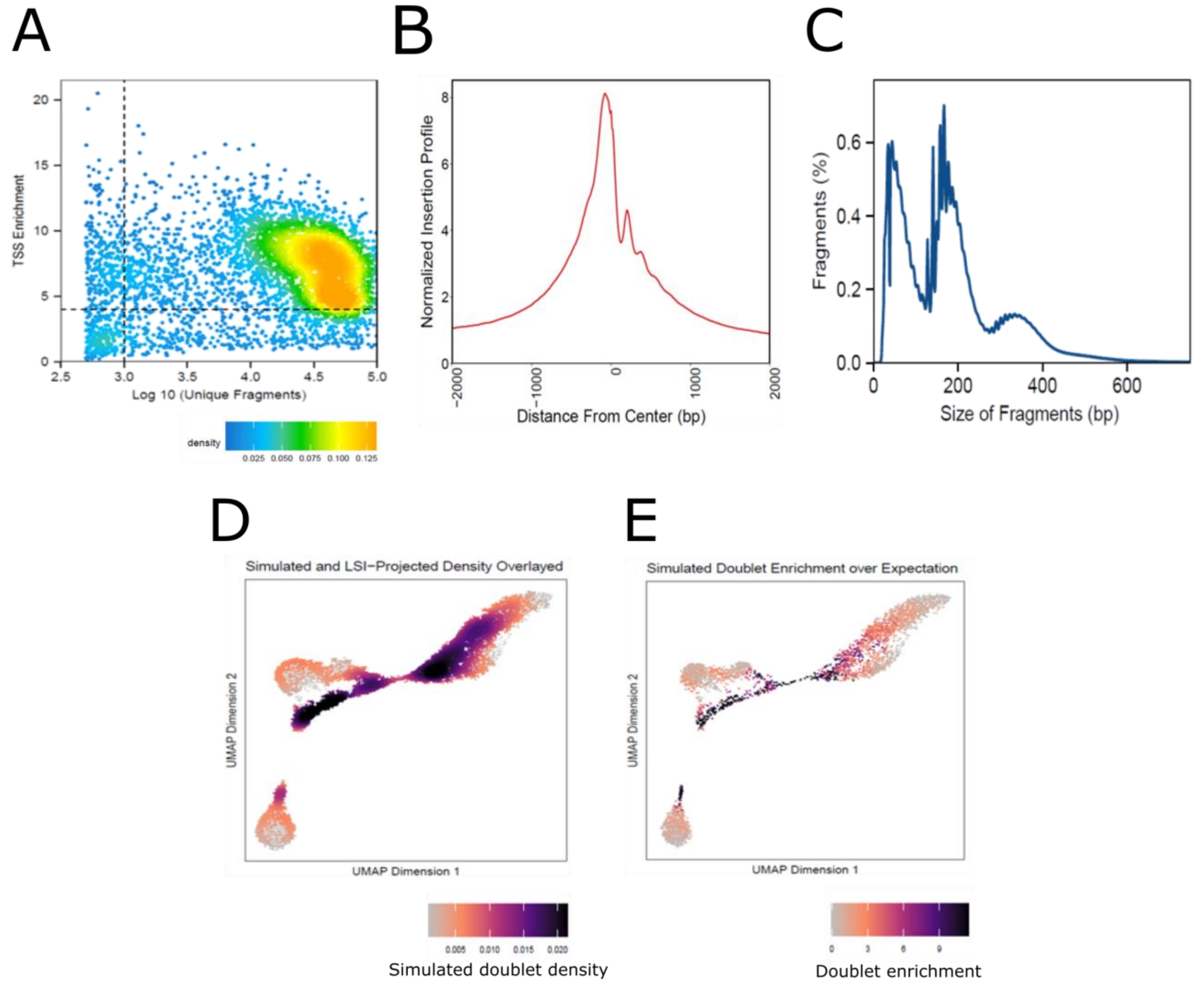


**Figure S9. Quality control & doublet identification for scATAC-seq profiles from primary tumor biopsy.** (A) QC filtering plots for the UW7 parental tumor samples used for scATAC-seq analysis. Transcription start site (TSS) enrichment score vs. log-transformed unique nuclear fragments per cell. Here TSS enrichment score is a calculated value representing the signal-to-background ratio. Colors indicate density of points in the plot. Of the total cells analyzed, 3,531 cells passed QC filters, located in the upper right quadrant. (B) TSS insertion profiles centered at all TSS regions. (C) Fragment size distributions for the UW7 primary tumor cells passing ArchR QC thresholds. UMAP plots of scATAC-seq data visualizing (D) simulated doublet density, (E) simulated doublet enrichment.

**Figure S10.**

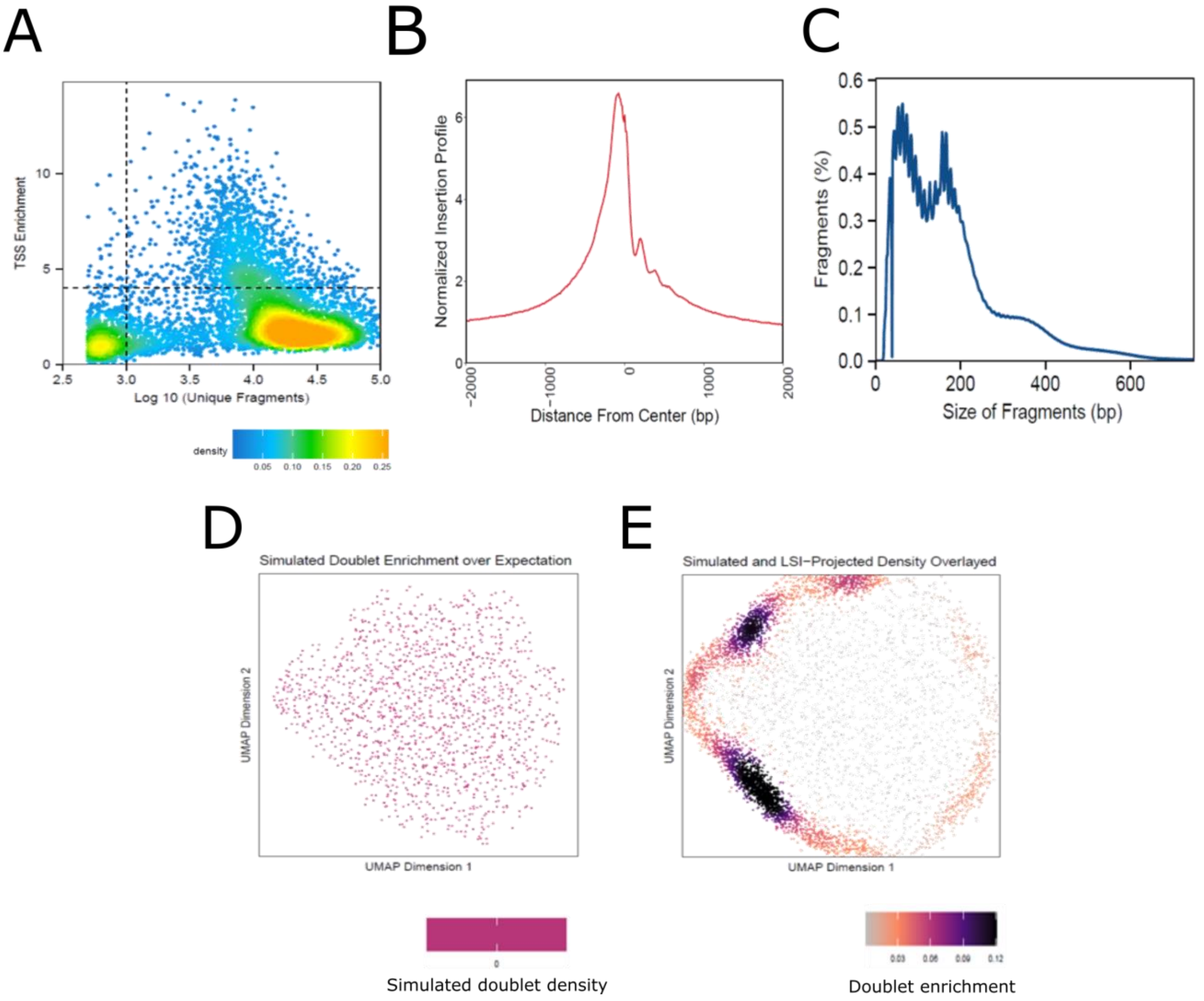


**Figure S10. Quality control & doublet identification for scATAC-seq profiles from recurrent tumor biopsy.** (A) QC filtering plots for the UW7 recurrent tumor used for scATAC-seq analysis. Transcription start site (TSS) enrichment score vs. log-transformed unique nuclear fragments per cell. Colors indicate density of points in the plot. Of the total cells analyzed, 1425 cells passed QC filters, located in the upper right quadrant. (B) TSS insertion profiles centered at all TSS regions. (C) Fragment size distributions for the UW7 primary tumor cells passing ArchR QC thresholds. UMAP plots of scATAC-seq data visualizing (D) simulated doublet density, (E) simulated doublet enrichment.
